# 19S proteasome loss causes monopolar spindles through ubiquitin-independent KIF11 degradation

**DOI:** 10.1101/2025.01.08.632038

**Authors:** Océane Marescal, Iain M. Cheeseman

**Affiliations:** Whitehead Institute for Biomedical Research, Cambridge, MA 02142; Department of Biology, Massachusetts Institute of Technology, Cambridge, MA 02142

## Abstract

To direct regulated protein degradation, the 26S proteasome recognizes ubiquitinated substrates through its 19S particle and then degrades them in the 20S enzymatic core. Despite this close interdependency between proteasome subunits, we demonstrate that knockouts from different proteasome subcomplexes result in distinct highly cellular phenotypes. In particular, depletion of 19S PSMD lid proteins, but not that of other proteasome subunits, prevents bipolar spindle assembly during mitosis, resulting in a mitotic arrest. We find that the monopolar spindle phenotype is caused by ubiquitin-independent proteasomal degradation of the motor protein KIF11 upon loss of 19S proteins. Thus, negative regulation of 20S-mediated proteasome degradation is essential for mitotic progression and 19S and 20S proteasome components can function independently outside of the canonical 26S structure. This work reveals a role for the proteasome in spindle formation and identifies the effects of ubiquitin-independent degradation on cell cycle control.

---

The ubiquitin-proteasome system is a central player in cellular regulation and is the major pathway for regulated protein degradation [1]. Proteins are marked for degradation by the covalent attachment of chains of ubiquitin to lysine residues, a post-translational modification known as poly-ubiquitination that targets a protein to the 26S proteasome [2, 3]. The 26S proteasome is a large protein complex whose subunits can be grouped into three subcomplexes [1, 4, 5] (Fig. 1A). The 20S core particle, composed of PSMA and PSMB subunits, is a barrel-shaped catalytic complex that contains the proteasome’s proteolytic activity [1, 4]. The 20S particle can be capped on either or both sides by the 19S regulatory particle, which can be further subdivided into the 19S base (PSMC/D subunits) and the 19S lid (PSMD subunits) [1, 4]. The unfolding and translocation of protein substrates into the core is driven by the AAA ATPase subunits of the 19S base, whereas the 19S lid is involved primarily in the recognition of ubiquitinated substrates and their de-ubiquitination [1, 3–8].

**Figure 1:**
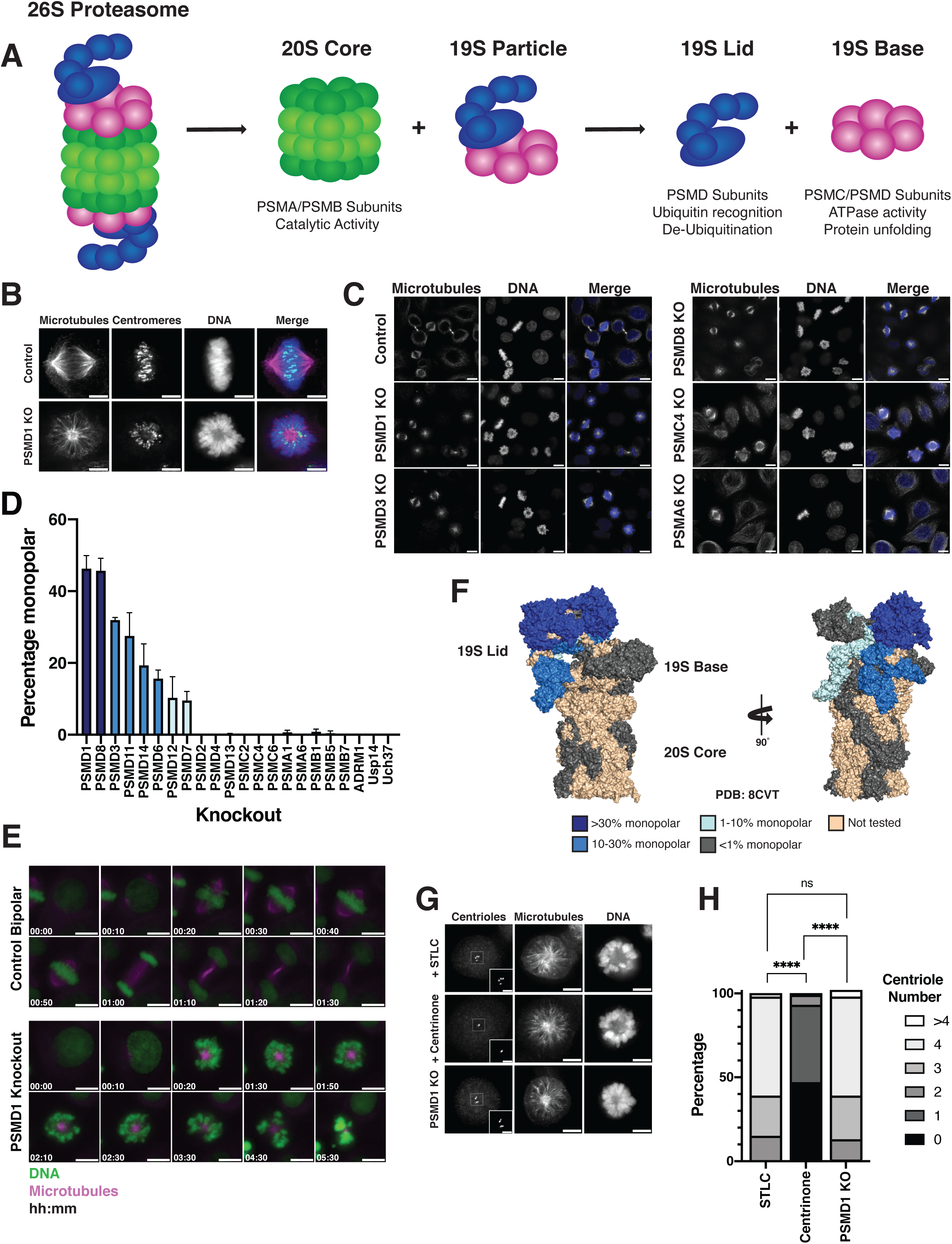
Depletion of a subset of 19S proteasome subunits induces monopolar spindles A. Diagram depicting 26S proteasome and its subcomplexes. B. Representative immunofluorescence images of mitotic cells with either a control bipolar spindle (top) or a monopolar spindle resulting from CRISPR-Cas9-based inducible knockout of PSMD1 (bottom). Scalebars = 5 µm. Images were acquired on different days and brightness is not scaled identically. C. Representative immunofluorescence images of asynchronous control HeLa cells, HeLa cells with inducible CRISPR-Cas9 knockout of PSMD1, PSMD3, or PSMD8, and HeLa cells transduced with lentivirus containing mCherry-expressing gene knockout plasmids for PSMC4 or PSMA6. Scalebars = 10 µm. Images were acquired on different days and brightness is not scaled identically. D. Graph showing the percentage of mitotic cells with monopolar spindles observed in cells with inducible CRISPR-Cas9 knockout of different proteasome components. Bars represent mean ± standard deviation of three or more replicates for each subunit. E. Live imaging stills of a PSMD1 inducible knockout cell entering mitosis or a control bipolar cell entering mitosis. Scalebars = 5 µm. Time units are hours:minutes. F. Structure of the 26S proteasome (PBD: 8CVT; [68]). Subunits are colored according to the percentage of monopolar cells observed in inducible CRISPR- Cas9 knockouts. G. Representative immunofluorescence images of monopolar mitotic cells from different conditions: cells treated with 10 µM STLC, cells treated with 200 nM centrinone [69], and inducible CRISPR-Cas9 PSMD1 knockout cells. STLC is a KIF11 inhibitor that does not affect centriole duplication [29], centrinone is a PLK4 inhibitor. Cells were stained for centrioles with Centrin 2 antibody. Regions enlarged in insets are outlined with dashed squares. Image brightness not identical. Scalebars for full-sized images = 5 µm. Scalebars for insets = 2 µm. H. Quantification of centriole number in different conditions: cells treated with 10 µM STLC, cells treated with 200 nM centrinone [69], and inducible CRISPR-Cas9 PSMD1 knockout cells. Bars show the percentage of cells in each centriole number category. Experiment was replicated 4 times, 36-51 cells were quantified for each condition for each replicate. P-values were calculated with chi-square tests. **** represents p< 0.0001, ns = not significant.

Ubiquitin-mediated degradation of specific substrates by the 26S proteasome plays a crucial role in the cell cycle and in mitotic progression [9–11]. For example, the degradation of cyclins by the ubiquitin pathway mediates each phase of the cell cycle [12], while anaphase onset during mitosis also relies on the ubiquitin-mediated degradation of specific protein components [13, 14]. In these cases, the ubiquitin system allows for the strict regulation of the timing, quantity, and specificity of protein degradation mediated by the intact 26S proteasome complex [9–11, 13, 14]. However, non-canonical proteasomal degradation, including ubiquitin-independent degradation or degradation by alternative proteasome complexes, can also occur [15–23], and their effects on the cell cycle are poorly understood. Identifying the consequences of such alternative modes of proteasome degradation on cell cycle and mitotic progression is integral for understanding both cell division and proteasome function.

A key step in mitosis is the formation of the mitotic spindle, a microtubule-based structure that binds to chromosomes during mitosis and segregates them into two daughter cells [24–26]. To achieve its function, the spindle must assemble into a bipolar configuration [24–26]. Bipolar spindle formation begins with the duplication of centrioles during interphase to form two distinct microtubule-organizing centers called centrosomes [24, 26, 27]. During mitosis, the centrosomes move to opposite ends of the cell to create two spindle poles from which microtubules emanate [24, 25]. These processes are tightly regulated and require multiple molecular players, including centriole duplication factors, microtubule-associated proteins, microtubule-nucleating components, motor proteins, and kinases [24–28]. Aberrant spindle assembly can lead to the formation of multipolar spindles with multiple spindle poles or monopolar spindles in which chromosomes surround a single pole [24, 27]. Such spindle assembly defects have deleterious consequences for the cell, resulting in mitotic arrest, cell death, or incorrect segregation of genetic material [27, 29].

Here, based on the analysis of cellular phenotypes, we uncover a novel role for proteasome 19S lid subunits in ensuring mitotic bipolar spindle assembly. Using systematic knockout of proteasome components, we tested the effects of individual proteasome subunit depletion on cellular phenotype. We find that the depletion of 19S lid proteins (PSMD subunits), but not that of other proteasome components, leads to the formation of monopolar spindles. We also show that this phenotype is caused by the ubiquitin-independent proteasomal degradation of the key spindle assembly factor, the kinesin motor protein KIF11, which occurs upon the loss of PSMD subunits from the proteasome core. Thus, the 19S lid restrains 20S proteasome degradation of specific cellular proteins, where the absence of negative regulation and changes in ubiquitin-independent degradation leads to failure in spindle assembly and mitotic arrest.

## Results

### Depletion of a subset of 19S proteasome subunits induces monopolar spindles

The role of the proteasome in cell cycle progression is typically regarded from the standpoint of ubiquitin-dependent protein degradation mediated by the entire 26S proteasome. However, this perspective fails to consider the potential for separable individual contributions of 19S and 20S subcomplexes to cellular function. As recent work has highlighted the possibility for increased separation of function of proteasome subcomplexes [19, 20, 30–32], we hypothesized that different proteasomal subunits could have unique roles in mitosis and cellular morphology. Therefore, to test the contributions of different proteasome subunits and their cellular phenotypes, we systematically generated individual CRISPR-Cas9-mediated knockouts of 19S and 20S proteasome genes using a doxycycline-inducible knockout system in HeLa cells (Table 1, [33]).

Knockout of a subset of PSMD subunits from the 19S particle resulted in a surprising mitotic phenotype: the formation of mitotically arrested cells with monopolar spindles, a spindle assembly defect where cells have only a single spindle pole instead of the normal bipolar spindle structure (Fig. 1 B, C, D, E). These subunits are primarily structural components located within the lid of the 19S complex, as well as the 19S base subunit PSMD1 (Fig. 1D, F). By contrast, knockout of PSMA or PSMB subunits from the 20S core particle or PSMC ATPase subunits did not produce any monopolar mitotic cells (Fig. 1C, D, F). We also used RNAi to deplete proteasome subunits. Depletion of the 20S catalytic subunit PSMB7 by RNAi did not result in cells with monopolar spindles, whereas cells treated with siRNAs targeting the 19S subunit PSMD1 displayed a high percentage of monopolar cells (74%) (Extended Data Fig. 1A, B, C). The knockout of PSMD1 in other cell types, including the non-small cell lung cancer cell line, A549, and the mouse fibroblast cell line, 3T3, also resulted in mitotically arrested cells with monopolar spindles (Extended Data Fig. 1D).

As the proteasome had not previously been implicated in mitotic spindle formation, we sought to understand the basis for the monopolar phenotype. We first considered other factors whose loss would lead to similar spindle defects, such as genes involved in centriole duplication, whose elimination also results in monopolar spindle formation [24, 30]. We tested whether defective centriole duplication could explain the monopolar spindle phenotype in PSMD knockout cells. Knockout or inhibition of known centriole replication factors, such as PLK4 or SAS6, resulted in monopolar spindles with reduced centriole numbers (Fig. 1G, H; Extended Data Fig. 1E, F). In contrast, PSMD1 knockout cells had normal centriole numbers (Fig. 1G, H; Extended Data Fig. 1E, F, [29]). Thus, the knockout of PSMD subunits induces monopolar spindle formation without disrupting centriole duplication. Together, these results suggest that bipolar spindle assembly requires proper proteasome function and that only a specific subregion of the proteasome, the 19S lid, plays a unique role in regulating spindle bipolarity.

### Decrease in ubiquitin-dependent degradation is not the cause of monopolarity

One of the primary roles of the 19S particle is to recognize ubiquitinated substrates for proteasomal degradation [1]. Given this role, we next tested the effects of PSMD subunit depletion on the degradation of ubiquitinated proteins. We conducted a cycloheximide chase assay and used the levels of MDM2, a ubiquitinated protein with high turnover [34, 35], as a reporter for ubiquitin-mediated protein degradation. In control cells, the inhibition of new protein synthesis led to the rapid loss of MDM2 (Fig. 2A; [35]). By contrast, cells treated with the proteasome inhibitor MG-132 maintained high levels of MDM2 over time (Fig. 2B). Similarly, the knockout of the lid subunits PSMD1, PSMD8, and PSMD11, as well as the core and base subunits PSMB7 and PSMC4, each showed decreased degradation of MDM2 (Fig. 2A, Extended Data Fig. 2A, B, C).

**Figure 2:**
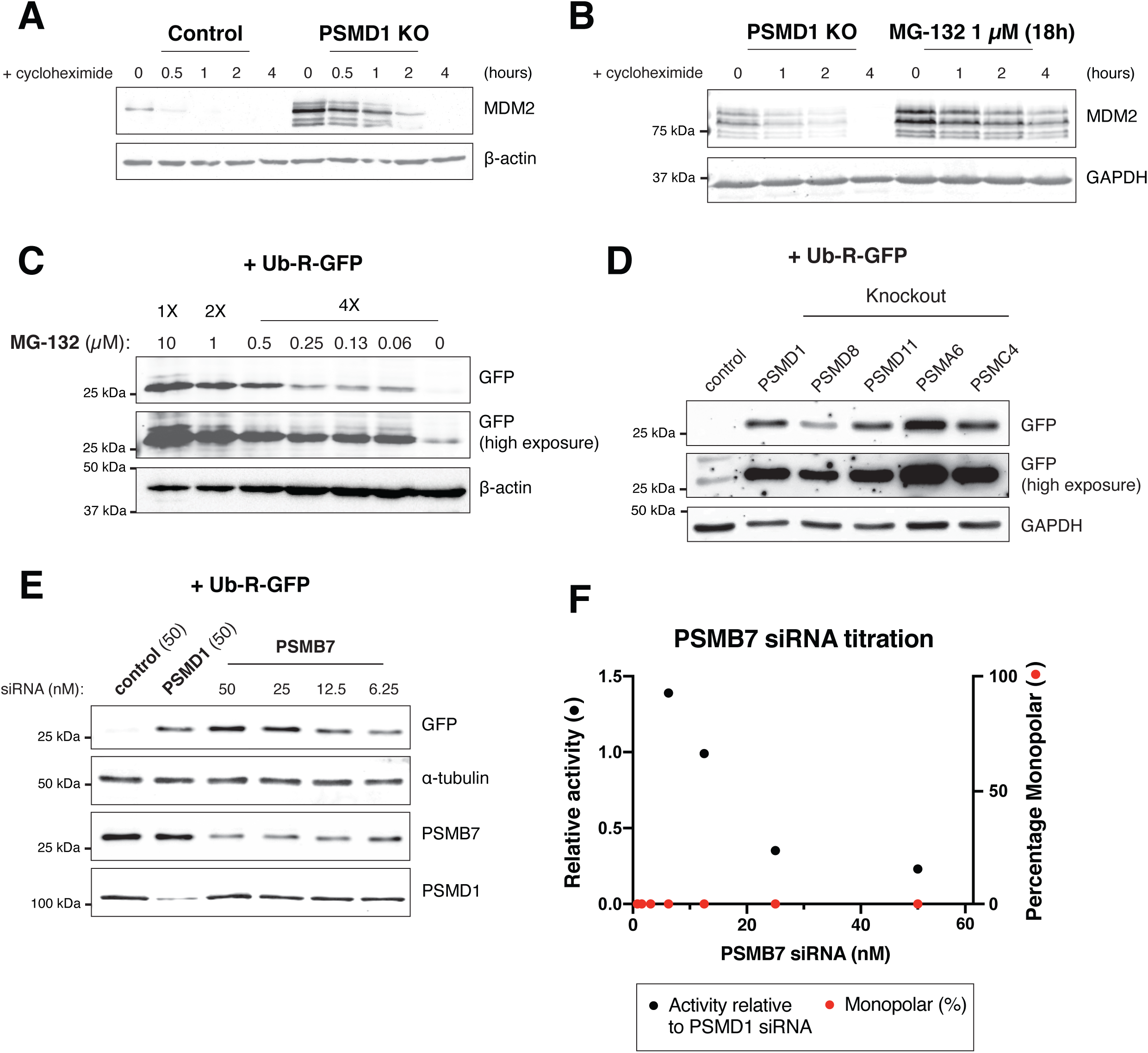
Decrease in ubiquitin-dependent degradation is not the cause of monopolarity A. Western blot of control or inducible CRISPR-Cas9 PSMD1 knockout cells treated with 50 µg/ml of cycloheximide for the indicated amount of time. Blot was incubated with MDM2 antibody. β-actin is used as a loading control. B. Western blot of inducible CRISPR-Cas9 PSMD1 knockout cells or cells treated with 1 µM MG-132 treated with 50 µg/ml of cycloheximide for the indicated amount of time. MG-132 was added to cells 18 hours prior to cycloheximide addition. Blot was incubated with MDM2 antibody. GAPDH is used as a loading control. C. Western blot of cells transfected with Ub-R-GFP construct treated with the indicated concentrations of MG-132. Ub-R-GFP was transfected 48 hours prior to collecting cells and MG-132 was added 24 hours prior to collecting cells. Blot was incubated with GFP antibody. Lower and higher exposure times are shown. β-actin is used as a loading control. D. Western blot of control cells or proteasome subunit knockout cells transfected with Ub-R-GFP construct. Knockouts were generated by transducing HeLa cells with lentivirus containing the mCherry-expressing gene knockout plasmids for the indicated genes. Cells were then sorted for mCherry positive cells by FACS 24 hours after transduction. 24 hours after sorting, cells were transfected with Ub-R- GFP plasmid and collected one day later. Blot was incubated with GFP antibody. Lower and higher exposure times are shown. GAPDH is used as a loading control. E. Western blot of cells treated with 50 nM control siRNAs, 50 nM PSMD1 siRNAs, and the indicated concentrations of PSMB7 siRNAs, transfected with Ub-R-GFP construct. Blot was incubated with GFP, PSMB7, or PSMD1 antibody. α-tubulin is used as a loading control. F. Graph showing the amount of proteasome activity in cells treated with different concentrations of PSMB7 siRNAs relative to the activity in cells treated with 50 nM PSMD1 siRNAs (black dots) and the percentage of monopolar cells observed by immunofluorescence in cells treated with different concentrations of PSMB7 siRNAs (red dots). Y-axis indicates the relative amount of proteasome activity or percentage of monopolar cells. Activity values were calculated by quantifying the intensity of GFP bands from (E). Each measurement for the PSMB7 knockdown cells was divided by the value measured for PSMD1 knockdown cells. Percentage of monopolar cells was zero for all concentrations over three replicates of the experiment.

We also assessed cellular degradation by measuring the stabilization of an exogenously expressed degron-tagged GFP construct (Ub-R-GFP; [36]). Cells treated with MG-132 showed an increase in the levels of Ub-R-GFP, indicating decreased degradation of the ubiquitinated construct (Fig. 2C, Extended Data Fig. 2D). Depletion of the 20S core components PSMA6 or PSMB7, or that of the ATPase PSMC4, also stabilized Ub-R-GFP (Fig. 2D, 2E; Extended Data Fig. 2F; Table 2). Similarly, knockouts of the 19S subunits PSMD1, PSMD8, and PSMD11 each stabilized the Ub-R-GFP fusion (Fig. 2D, 2E; Extended Data Fig. 2E; Table 2). We also observed a global increase in the total levels of ubiquitin-conjugated proteins in cells treated with PSMD1 siRNAs, PSMB7 siRNAs, or MG-132 (Extended Data Fig. 2G). Thus, although only the knockout of 19S lid (PSMD) subunits induces monopolarity, depletion of components from all sub-regions of the proteasome decreases ubiquitin-mediated degradation, consistent with the previously established function for these proteasome components.

Although the depletion of PSMD subunits reduced ubiquitin-mediated degradation, in some cases this reduction was less potent than that observed for 20S core subunit-depleted cells. Thus, we wondered whether a partial reduction of proteasome activity in PSMD knockouts could be the cause of monopolar spindle formation. To test this, we titrated siRNAs against PSMB7, a 20S core component, to induce varying levels of cellular protein degradation. Lower concentrations of PSMB7 siRNAs resulted in higher proteasome activity as measured by the degradation of Ub-R- GFP (Fig. 2E). However, we did not observe monopolar cells at any concentration of PSMB7 siRNAs, even at concentrations where proteasome activity was equivalent to that found in cells treated with PSMD1 siRNAs (12.5 nM PSMB7 siRNA) (Fig. 2E, F). Thus, the knockout of components from any subcomplex of the proteasome decreases ubiquitin-mediated degradation, but the relative decrease in cellular degradation does not account for the cellular monopolar spindle phenotype observed in PSMD-depleted cells.

### Eg5/KIF11 is uniquely lost in PSMD-depleted cells

PSMD knockout cells have a potent monopolar spindle phenotype that could not be explained by defects in centriole duplication (Fig. 1G, H; Extended Data Fig. 1E, F). To determine what other factors were responsible for the observed monopolarity in PSMD- depleted cells, we compared global changes in protein abundances between cells treated with either control siRNAs or PSMD1 siRNAs using Tandem Mass Tag (TMT) quantitative mass spectrometry (Fig. 3A; Extended Data Fig. 3A; Supplementary Table 1, 2). Given the decrease of ubiquitin-mediated degradation in PSMD1-depleted cells (Fig. 2A, D, E; Extended Data Fig. 2E), we first searched for an upregulated protein whose lack of degradation could lead to monopolar spindle formation. Although several proteins that undergo ubiquitin-mediated degradation were increased in PSMD1 knockdown cells (Supplementary Table 1, 2, [12, 37–43]), none were relevant to spindle assembly. However, in both interphase and mitotic cells, one of the most significantly *downregulated* factors in PSMD1-depleted cells was the protein Eg5/KIF11 (Fig. 3A, Extended Data Fig. 3A, Supplementary Table 1, 2). KIF11 is a kinesin-5 motor protein that separates spindle poles during spindle assembly [24–26, 44, 45]. Loss of KIF11 activity through depletion or chemical inhibition results in potent monopolar spindle formation (Fig. 3B, [24, 25, 44–46]).

**Figure 3:**
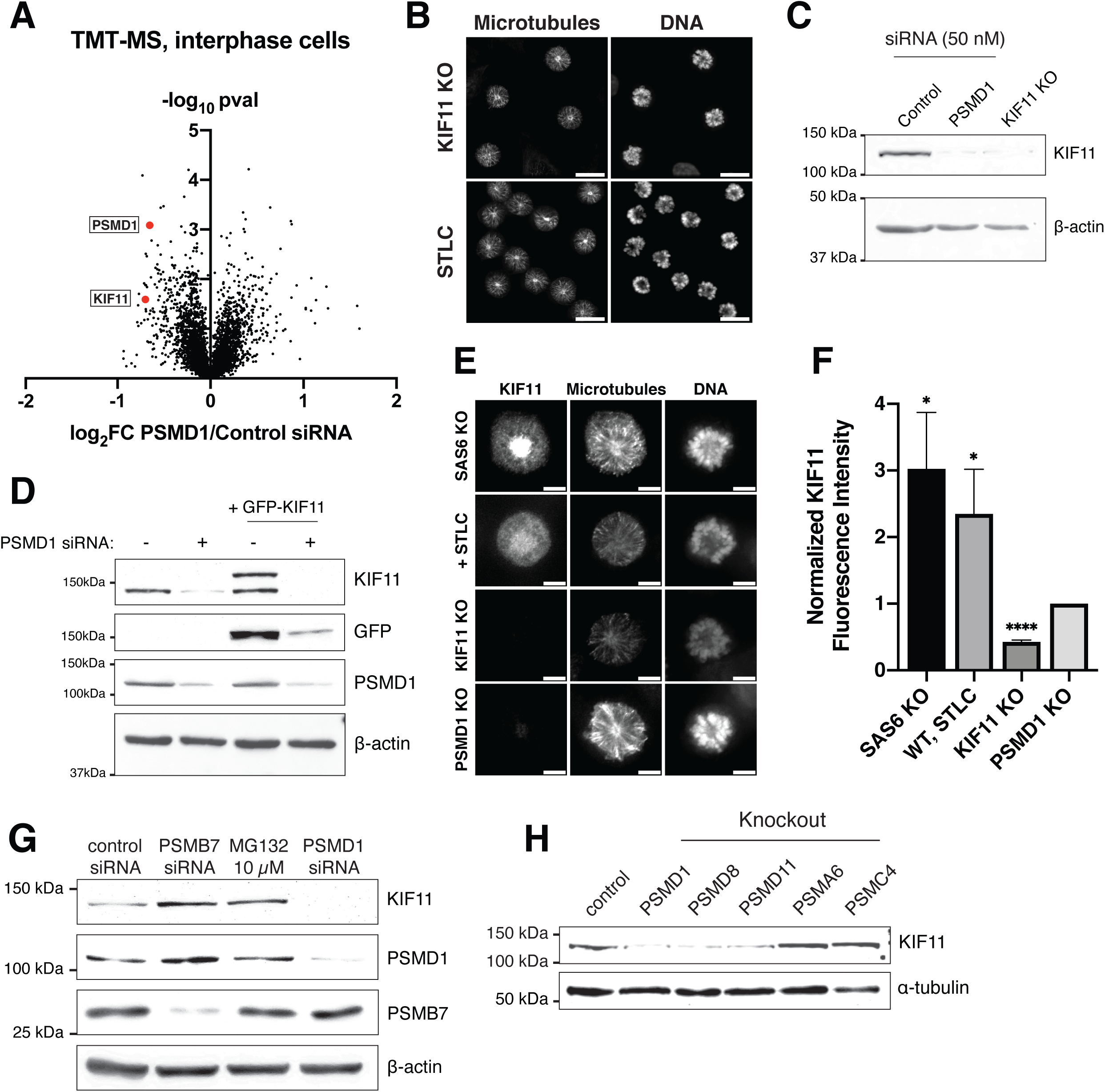
Eg5/KIF11 is uniquely lost in PSMD-depleted cells A. Comparison of protein abundances in cells treated with 50 nM PSMD1 siRNAs and cells treated with 50 nM control siRNAs. Cells were treated with siRNAs for 48 hours and arrested in S phase with 2 mM thymidine prior to harvesting. Protein abundances were obtained using TMT-based (Tandem Mass Tag) quantitative mass spectrometry of three replicates for each condition. Volcano plot shows log_2_ of protein abundances in PSMD1 knockdown cells divided by protein abundances in control knockdown cells vs. -log_10_ of p-values. Values on the right of the plot represent proteins with increased abundance in PSMD1 knockdown cells and values on the left represent proteins with decreased abundance in PSMD1 knockdown cells. PSMD1 and KIF11 are highlighted in red. B. Representative immunofluorescence images of KIF11 CRISPR-Cas9 inducible knockout cells and control cells treated with 10 µM *S*-Trityl-L-cysteine (STLC). Scalebars = 20 µm. Cells were fixed and stained on different days and brightness is not scaled identically. C. Western blot of cells treated with control siRNAs, cells treated with PSMD1 siRNAs, and inducible CRISPR-Cas9 KIF11 knockout cells. Blot was incubated with KIF11 antibody. β-actin is used as a loading control. D. Western blots of control or GFP-KIF11-expressing cell line treated with 50 nM control (-) or PSMD1 (+) siRNAs. Blots were incubated with KIF11, GFP, and PSMD1 antibodies. Separate blots were used for each antibody with the same samples and same amounts loaded. β-actin is used as a loading control. E. Representative immunofluorescent images of monopolar mitotic cells showing KIF11 levels in different conditions: SAS6 inducible CRISPR-Cas9 knockout (positive control), control cells treated with 10 µM *S*-Trityl-L-cysteine (STLC), KIF11 inducible CRISPR-Cas9 knockout, and PSMD1 inducible CRISPR-Cas9 knockout. Scalebar = 5 µm. F. Quantification of KIF11 fluorescence intensity in monopolar mitotic cells in different conditions. Fluorescence intensity values were normalized to levels in PSMD1 knockout cells. Bars represent mean ± standard deviation. P-values were calculated with two-tailed Welch’s t-tests comparing each condition to PSMD1 knockout: p = 0.0275 for control + STLC, p = 0.0173 for SAS6 knockout, and p < 0.0001 for KIF11 knockout. * represents p < 0.05 and **** represents p < 0.0001. Experiment was replicated 4 times; 29-92 cells were quantified for each condition for each replicate. G. Western blots of cells treated with either 50 nM control siRNAs, 50 nM PSMB7 siRNAs, 50 nM PSMD1 siRNAs, or 10 µM MG132. Blots were incubated with KIF11 antibody, PSMD1 antibody, and PSMB7 antibody. β-actin is used as a loading control. Separate blots were used for each antibody with the same samples and same amounts loaded. β-actin is shown for KIF11 blot but loading control was also conducted for PSMD1 and PSMB7 blot with similar results. H. Western blot of cells transduced with lentivirus containing the mCherry- expressing gene knockout plasmids for PSMD1, PSMD8, PSMD11, PSMA6, or PSMC4. Cells were sorted for mCherry. Control cells are untransduced parental cells. Blot was incubated with KIF11 antibody. α-tubulin is used as a loading control.

Consistent with the reduction in KIF11 protein levels observed by mass spectrometry, we observed the loss of the endogenous KIF11 protein in PSMD1 knockdown cells using western blotting (Fig. 3C). Similarly, we found that exogenously expressed GFP-tagged KIF11 was also lost in PSMD1 knockdown cells (Fig. 3D). In addition, we detected decreased KIF11 spindle localization in PSMD1, PSMD3, PSMD8, and PSMD11 knockout cells based on immunofluorescence against the endogenous protein (Fig 3E, F; Extended Data Fig. 3B, C). In contrast, depletion of PSMA6, PSMB7, or PSMC4, or the addition of MG-132, did not decrease KIF11 levels (Fig 3G, H) and instead resulted in an accumulation of KIF11. Thus, although depletion of other proteasome subunits or addition of a chemical proteasome inhibitor increases KIF11 levels, knockout of PSMD subunits leads to a unique loss of KIF11 protein.

### Loss of KIF11 contributes to the monopolar spindle phenotype in PSMD-depleted cells

We next evaluated whether the loss of KIF11 is sufficient to explain spindle monopolarity in PSMD-depleted cells. Ectopically expressed GFP-KIF11 was also degraded in PSMD1 knockdown cells (Fig. 3D) and thus failed to restore bipolar spindle formation (Fig. 6E, F). We therefore used alternate methods to evaluate the contributions of KIF11. We predicted that a reduction of KIF11 levels in PSMD-depleted cells would sensitize these cells to further KIF11 inhibition. We therefore analyzed the synergistic phenotypes between partial PSMD1 depletion using RNAi and partial inhibition of KIF11 activity using the chemical inhibitor *S*-Trityl-L-cysteine (STLC; [29]). Although treating cells with low concentrations of STLC or low concentrations of PSMD1 siRNAs alone did not result in substantial monopolarity (Fig. 4A, Extended Data Fig. 4A, B, C), simultaneously treating cells with both STLC and PSMD1 siRNAs resulted in an almost 3-fold increase in monopolar cells (Fig. 4A; Extended Data Fig. 4C).

**Figure 4:**
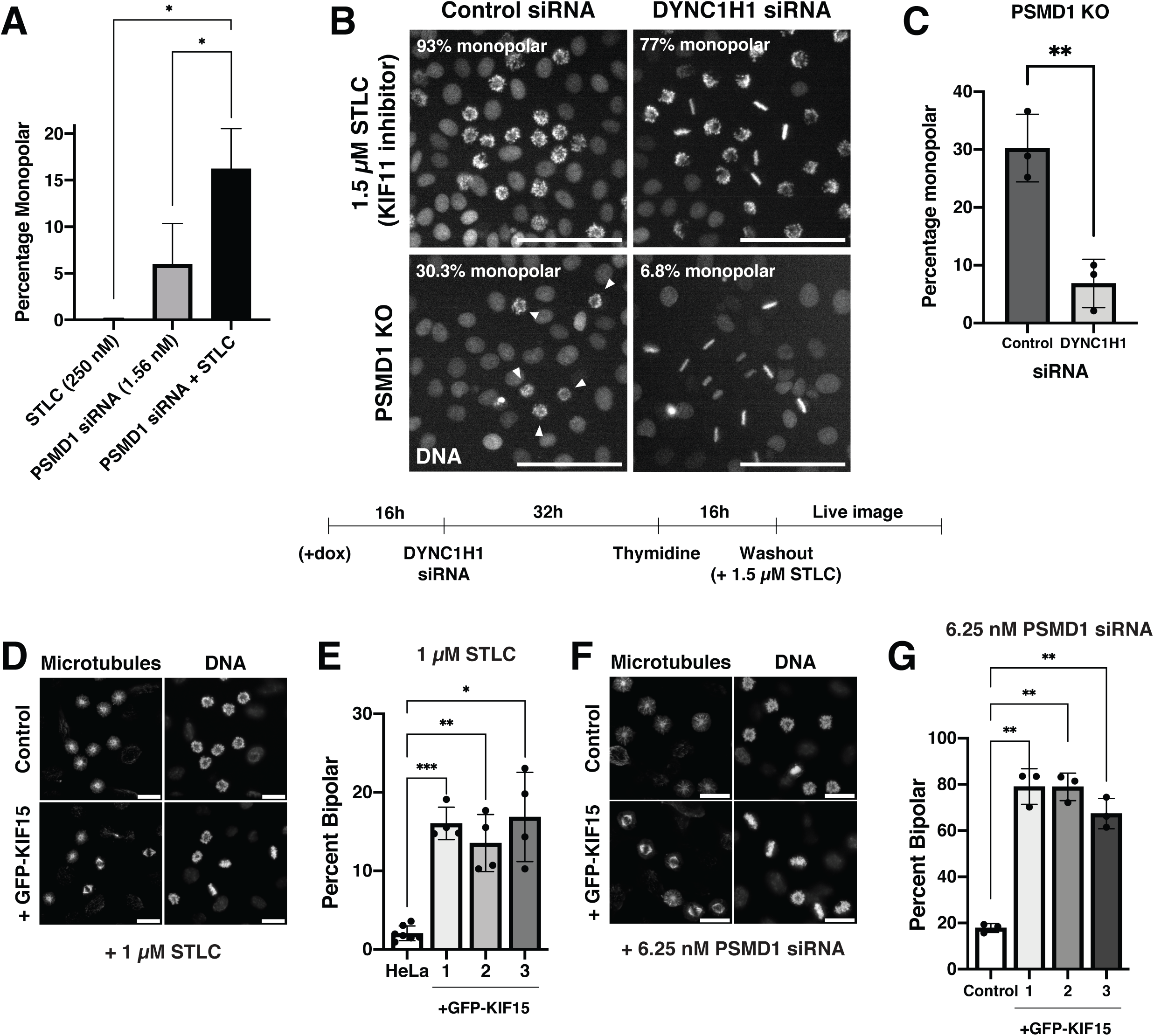
Loss of KIF11 contributes to the monopolar spindle phenotype in PSMD-depleted cells A. Quantification of live imaging synergy experiment. Bar graph plots the percentage of cells entering mitosis with monopolar spindles in cells treated with low dose (250 nM) STLC alone, low dose (1.56 nM) PSMD1 siRNAs alone, or both 250 nM STLC and 1.56 nM PSMD1 siRNAs together. P-values were calculated with two-tailed Welch’s t-tests, p = 0.023 for STLC vs. siRNA + STLC and p = 0.045 for PSMD1 siRNA vs. siRNA + STLC. * represents p < 0.05. Experiment was replicated 3 times and between 1090-2527 mitotic cells were counted for each condition for each replicate. B. Representative stills of live imaging experiments. *Top Panels*: Cells were treated with either 50 nM control siRNAs or 50 nM Dynein Heavy Chain siRNAs (DYNC1H1) and then with 1.5 µM of the KIF11 inhibitor, STLC. Experiment was conducted once, as these results have been previously studied [48]. *Bottom Panels*: Doxycycline was added to cells to induce knockout of PSMD1. Cells were then treated with either 50 nM control siRNAs or 50 nM Dynein Heavy Chain siRNAs (DYNC1H1). White arrows point towards monopolar cells. Scalebars = 100 µm. A diagram of the experimental timeline is displayed underneath the images. C. Quantification of dynein rescue experiment from Fig. 4B. Bar graph shows the percentage of cells entering mitosis with monopolar spindles in PSMD1 knockout cells treated with 50 nM of either control or dynein heavy chain siRNAs. P- values were calculated with two-tailed Welch’s t-tests, p = 0.0064. ** represents p < 0.01. Experiment was conducted 3 times and between 1215-1521 mitotic cells were counted for each condition in each replicate. D. Representative immunofluorescence images of either control or GFP-KIF15- expressing cells treated with 1 µM STLC. Scalebars = 20 µm. Image brightness is not scaled identically. E. Bar graph showing the percentage of mitotic cells with bipolar spindles in control HeLa cells or 3 different clones of a GFP-KIF15-overexpressing cell line treated with 1 µM STLC. P-values were calculated with two-tailed Welch’s t-tests, p = 0.0003 between HeLa and clone 1, p = 0.0068 between HeLa and clone 2, p =0.013 between HeLa and clone 3. *** represents p < 0.001, ** represents p < 0.01, * represents p < 0.05. Experiments were replicated 3 times and between 208-555 mitotic cells were counted for each condition in each replicate. F. Representative immunofluorescence images of either control or GFP-KIF15- expressing cells treated with 6.25 nM PSMD1 siRNAs. Scalebars = 20 µm. G. Bar graph showing the percentage of mitotic cells with bipolar spindles in control HeLa cells or 3 different clones of a GFP-KIF15-overexpressing cell line treated with 6.25 nM PSMD1 siRNAs. P-values were calculated with two-tailed Welch’s t-tests, p = 0.0035 between HeLa and clone 1, p = 0.0015 between HeLa and clone 2, p = 0.0034 between HeLa and clone 3. ** represents p < 0.01. Experiments were replicated 3 times and between 121-318 mitotic cells were counted for each condition in each replicate.

KIF11 separates spindle poles by producing outward force on the spindle [47]. Thus, if the loss of KIF11 contributes to spindle monopolarity in PSMD-depleted cells, either decreasing inward force or restoring outward force on the spindle should rescue spindle bipolarity. Prior work found that the depletion of the motor protein dynein rescues bipolar spindle formation in cells with reduced KIF11 activity by eliminating opposing inward force on the spindle (Fig. 4B *Top Panel*; [25, 48]). To test whether dynein depletion can also rescue monopolarity induced by loss of PSMD1, we treated PSMD1 knockout cells with siRNAs against dynein heavy chain (DYNC1H1) (Extended Data Fig. 4D). PSMD1 knockout cells treated with DYNC1H1 siRNAs formed significantly fewer monopolar spindles than those treated with control siRNAs (Fig. 4B *Bottom Panel*, C). Similarly, siRNAs against the dynein-binding protein Lis1 also rescued spindle bipolarity in PSMD1 knockouts (Extended Data Fig. 4E).

Reciprocally, restoring outward spindle forces in cells with low KIF11 levels is predicted to rescue bipolar spindle formation. Previous work found that the motor protein KIF15 (kinesin-12) acts in overlapping pathways with KIF11 to push spindle poles apart [49]. The overexpression of KIF15 can rescue the loss of KIF11 activity [25, 46], such that GFP-KIF15 overexpression restored spindle bipolarity in cells treated with STLC (Fig. 4D, E [46]; Extended Data Fig. 4F, G). Similarly, in cells treated with PSMD1 siRNAs, we found that cells overexpressing GFP-KIF15 had a significantly higher percentage of bipolar cells than control cells (Fig. 4F, G). Thus, KIF15 overexpression can rescue the effects PSMD1 depletion, despite there being no difference in KIF15 protein levels in PSMD1 knockdown cells (Extended Data Fig. 4H).

Thus, experimental manipulations known to rescue the loss of KIF11 activity, by either increasing the amount of outward force (KIF15 overexpression) or decreasing the amount of inward force on the spindle (Dynein Heavy Chain or Lis1 RNAi), rescue PSMD subunit depletion. Together, these data suggest that the loss of KIF11 in PSMD- depleted cells contributes to the monopolar phenotype generated upon proteasome lid subunit depletion.

### KIF11 loss is mediated by ubiquitin-independent proteasomal degradation

We next sought to uncover the mechanism by which KIF11 is lost in PSMD-depleted cells. Although we found that the levels of the endogenous KIF11 mRNA were decreased upon PSMD1 knockdown, KIF11 mRNA levels also decreased significantly in PSMB7 knockdown cells, which do not have decreased KIF11 protein levels (Extended Data Fig. 5A, Fig. 3G). In addition, GFP-KIF11 protein expressed from an ectopic promoter was also lost in PSMD1 knockdown cells (Fig. 3D), despite there being no difference in the change in mRNA levels upon PSMD1 knockdown of the GFP- KIF11 construct relative to that of GFP alone (Extended Data Fig. 5B, C). These experiments suggest that KIF11 loss occurs at the protein level.

To test whether loss of KIF11 protein is mediated by the proteasome, we simultaneously depleted both PSMD1 and the 20S catalytic core component PSMB7 using RNAi. Although treating cells with PSMD1 siRNAs alone led to an almost complete loss of KIF11 protein, simultaneously depleting both PSMD1 and PSMB7 prevented the degradation of KIF11 (Fig. 5A) and led to a 4-fold decrease in the fraction of monopolar cells (Fig. 5B, C). Similarly, PSMA6 knockout cells treated with PSMD1 siRNAs retained KIF11 protein and showed a rescue of bipolar spindle formation (Extended Data Fig. 5D, E, F). Finally, addition of the chemical proteasome inhibitor MG-132 impeded degradation of KIF11 (Fig. 5D). Thus, genetic or chemical inhibition of proteasome activity rescues KIF11 protein levels and bipolar spindle formation in PSMD-depleted cells, suggesting that KIF11 is degraded by the proteasome despite an overall decrease in ubiquitin-mediated degradation (Fig. 2).

**Figure 5:**
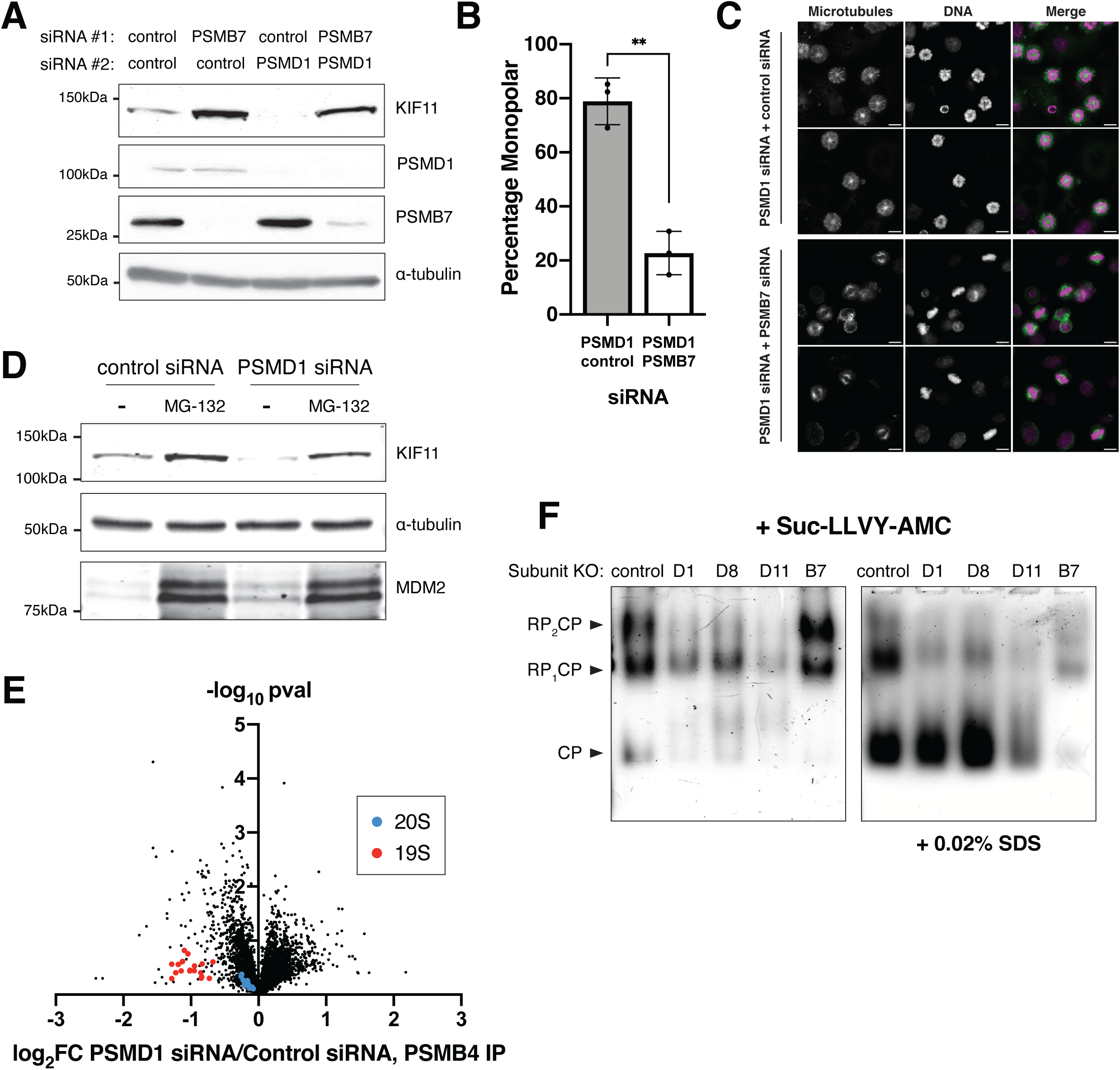
KIF11 loss is mediated by proteasomal degradation A. Western blots of double knockdown cells. Cells were treated with 50 nM of the indicated siRNA #1 for 72 hours and the indicated siRNA #2 for 48 hours before collection. Blots were incubated with KIF11, PSMD1, PSMB7 antibodies. α- tubulin is used as a loading control. Separate blots were used with the same samples and same amounts loaded. One blot was incubated with PSMB7/D1 antibodies and the other with KIF11/α-tubulin antibodies. B. Quantification of immunofluorescence experiment from PSMD1/PSMB7 double knockdown cells. Bar graph shows the percentage of mitotic cells with monopolar spindles for cells treated with both 12.5 nM PSMD1 and 50 nM control siRNAs or cells treated with 12.5 nM PSMD1 and 50 nM PSMB7 siRNAs. P-value was calculated with two-tailed Welch’s t-tests, p = 0.0012. ** represents p < 0.01. Experiment was conducted 3 times. C. Representative immunofluorescence images for double knockdown experiment from 5B. Top two panels show HeLa cells treated with PSMD1 siRNAs and control siRNAs. Bottom two panels show HeLa cells treated with PSMD1 siRNAs and PSMB7 siRNAs. Scale bars = 10 µm. Image brightness not scaled the same. D. Western blots of cells treated with 50 nM of the indicated siRNAs with or without 10 µM MG-132. siRNAs were added 48 hours prior to collection and MG-132 was added 24 hours prior to collection. Blots were incubated with KIF11 and MDM2 antibodies. α-tubulin is used as a loading control. Separate blots were used with the same samples and same amounts loaded. One blot was incubated with KIF11 and α-tubulin antibodies and the other with MDM2 antibody. E. Comparison of protein abundances of GFP immunoprecipitations from GFP- PSMB4 cells treated with 50 nM PSMD1 siRNAs and cells treated with 50 nM control siRNAs. Cells were treated with siRNAs for 48 hours and with 2 mM Thymidine 24 hours prior to harvesting. Protein abundances were obtained using TMT-based (Tandem Mass Tag) quantitative mass spectrometry of three replicates for PSMD1 RNAi condition and two replicates for control RNAi condition. Volcano plot shows log_2_ of protein abundances in PSMD1 knockdown cells divided by protein abundances in control knockdown cells vs. -log_10_ of p- values. 19S components and 20S components are highlighted in red and blue, respectively. F. Native gel followed by Suc-LLVY-AMC assay using different proteasome inducible knockout cell lines. Knockout of indicated proteasome components was induced 3-4 days prior to collecting cells. Native gel and incubation in Suc-LLVY- AMC was performed as described in the Methods. Bands corresponding to double capped (RP_2_CP), single capped (RP_1_CP), or uncapped 20S proteasomes (CP) are indicated. Addition of 0.02% SDS opens 20S proteasome substrate channel and allows for detection of 20S proteasome activity.

**Figure 6:**
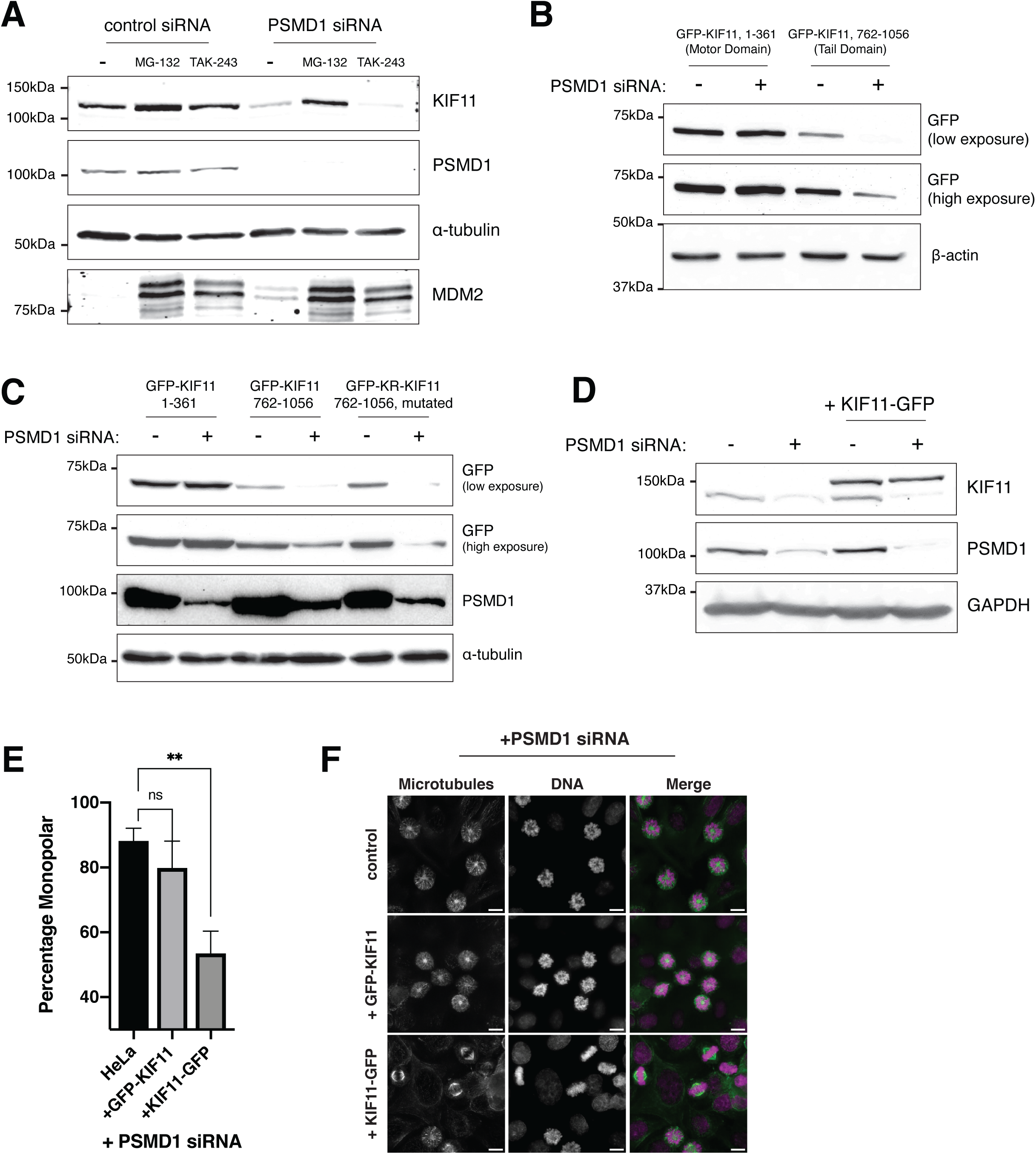
Ubiquitin-independent proteasomal degradation of KIF11 is mediated by its unstructured C-terminus A. Western blots of cells treated with 50 nM of the indicated siRNAs and the indicated compounds. MG-132 was used at 10 µM and TAK-243 at 300 nM. siRNAs were added 48 hours prior to collection and compounds were added 24 hours prior to collection. Blots were incubated with KIF11, PSMD1, and MDM2 antibodies. α-tubulin is used as a loading control. Separate blots were used with the same samples and same amounts loaded. One blot was incubated with KIF11, PSMD1, and α-tubulin antibodies and the other with MDM2 antibody. B. Western blot of cell lines expressing the indicated GFP-KIF11 truncation constructs treated with 50 nM control (-) or PSMD1 (+) siRNAs. Blot was incubated with GFP antibody. Lower and higher exposure times are shown. β- actin is used as a loading control. C. Western blot of cell lines expressing the indicated GFP-KIF11 truncation constructs treated with 50 nM control (-) or PSMD1 (+) siRNAs. Blot was incubated with GFP and PSMD1 antibodies. Lower and higher exposure times are shown. α-tubulin is used as a loading control. D. Western blot of control cells or cell line expressing C-terminally-tagged KIF11- GFP construct. Cells were treated with either 50 nM control (-) or PSMD1 (+) siRNAs. Blot was incubated with KIF11 and PSMD1 antibodies. GAPDH is used as a loading control. E. Quantification of immunofluorescence experiment for cells expressing KIF11 constructs. Bar graph shows the percentage of mitotic cells with monopolar spindles for parental HeLa cells, a cell line expressing N-terminally-tagged GFP- KIF11, and a cell line expressing C-terminally-tagged KIF11-GFP treated with 12.5 nM PSMD1 siRNAs. ** indicates p < 0.01, ns = not significant. F. Representative immunofluorescence images from 5H. Scale bar = 10 µm.

Depletion of 19S lid subunits is predicted to prevent formation of a capped 26S proteasome but could potentially leave the 20S catalytic core intact [50–53]. We therefore measured the association of 19S subunits to the 20S core using immunoprecipitation followed by quantitative mass spectrometry (IP-MS) (Fig. 5E, Supplementary Table 3). Immunoprecipitation of the core subunit PSMB4 in PSMD1 knockdown cells pulled down substantially less of each 19S subunit when compared to control knockdown cells, suggesting a dissociation of the proteasome 19S particle from the core upon PSMD1 knockdown (Fig. 5E). Conversely, the interaction of PSMB4 with other core subunits was unaffected (Fig. 5E, Supplementary Table 3). We also used native gels to measure the ratio of different proteasome complexes and their catalytic activity (Fig. 5F). Native gels separate whole proteasome complexes by size without compromising their activity, allowing for the *in vitro* cleavage of the fluorescent substrate Suc-LLVY-AMC (Suc-Leu-Leu-Val-Tyr-AMC; [54–56]) to detect which populations of proteasome complexes are capable of substrate cleavage. We observed a decrease in the 26:20S proteasome ratio in PSMD knockout cells but found that the accumulated 20S cores retained their catalytic cleavage activity (Fig. 5F), consistent with decreased ubiquitin-mediated degradation but preservation of 20S core proteasome activity.

Although poly-ubiquitination and subsequent degradation by the 26S proteasome is the most frequently used cellular degradation pathway, some substrates have also been shown be degraded in a ubiquitin-independent manner [15–17]. As the 19S lid is the ubiquitin-recognizing component of the 26S proteasome and dissociates from the 20S core upon PSMD depletion, we hypothesized that the degradation of KIF11 in lid- depleted cells could be proteasome-dependent but ubiquitin-independent. To test whether degradation of KIF11 is ubiquitin-independent, we treated PSMD1 knockdown cells with either MG-132 to inhibit proteasome catalytic activity or TAK-243 to inhibit ubiquitination. TAK-243 (MLN7243) is an inhibitor of the ubiquitin activating E1 enzyme UBA1 and disrupts ubiquitination upstream of proteasome activity [57]. Although addition of either MG-132 or TAK-243 decreased degradation of the ubiquitin-dependent substrate MDM2, only the addition of MG-132 rescued KIF11 levels in PSMD1-depleted cells (Fig. 6A). Thus, TAK-243-treated PSMD1 knockdown cells continue to degrade KIF11, even in the absence of E1 activity and ubiquitination. In contrast to PSMD1 knockdown cells, control cells treated with either MG-132 or TAK243 both accumulated KIF11 (Extended Data Fig. 5G).

To determine the region of KIF11 responsible for its degradation, we created truncations of the protein and assessed their degradation. In contrast to the full-length protein, the N-terminal motor domain of KIF11 (amino acids 1-361) did not show reduced protein levels upon PSMD1 knockdown (Fig. 6B). However, the depletion of PSMD1 *did* induce the loss of the KIF11 C-terminal tail domain (amino acids 762-1056) (Fig. 6B). Because ubiquitination of substrate proteins occurs through the covalent attachment of ubiquitin to lysine residues, we also created a lysine to arginine mutant of the C-terminal tail domain of KIF11 (GFP-KR-KIF11, 762-1056). Although the lysine-less KIF11 fragment would be incapable of being ubiquitinated, it was still degraded upon PSMD1 knockdown (Fig. 6C). Thus, the degradation of KIF11 depends on its C-terminal domain but not on its ubiquitination.

Previous cases of ubiquitin-independent degradation have been found to require the presence of an unstructured region to initiate translocation into the proteasome [18, 19, 58, 59]. Structural models of KIF11 predict the presence of a ∼130 amino acid intrinsically disordered region at the C-terminus of the protein (Extended Data Fig. 6A, 6B). Because our C-terminal construct was degraded in our truncation experiments (Fig. 6B, C), we tested whether this exposed disordered region was essential for the ubiquitin-independent degradation of KIF11. To do so, we tagged KIF11 with GFP at its C-terminus (KIF11-GFP). Although an N-terminally-tagged GFP-KIF11, whose C-terminus remains available, was degraded efficiently upon addition of PSMD1 siRNAs (Fig. 3D), we found that the C-terminally-tagged KIF11-GFP was not (Fig. 6D, Extended Data Fig. 6C). Correspondingly, we observed an increase in spindle bipolarity in cells treated with PSMD1 siRNAs expressing KIF11-GFP, but not in those expressing GFP- KIF11 (Fig. 6E, F). Thus, blocking KIF11’s free disordered C-terminal region with GFP prevented the protein’s degradation, and the consequent rescue of KIF11 levels was sufficient to rescue monopolar spindles induced by PSMD1 depletion.

Interestingly, structural predictions of KIF11 from other species suggest that the presence of this C-terminal disordered region is conserved in eukaryotes from fungi to vertebrates (Extended Data Fig. 6D). This conservation could indicate a drive to preserve KIF11 degradation through a ubiquitin-independent mechanism under conditions where a cell could tune its proteasome complex composition to regulate KIF11 abundance and spindle assembly.

From these data, we conclude that the loss of 19S lid components triggers the ubiquitin-independent proteasomal degradation of KIF11, which is dependent on its disordered C-terminus.

## Discussion

The canonical pathway for protein degradation is the ubiquitination of target proteins followed by their degradation by the 26S proteasome. Here we show that the depletion of different proteasome subunits can give rise to vastly different cellular behaviors, highlighting the capacity for independent functions of the 19S and 20S particles outside of the canonical 26S complex. We find that the knockout of 19S PSMD proteasome components causes a failure in spindle pole separation during mitosis, resulting in the formation of aberrant monopolar spindles. However, despite functioning in the same complex, other subunits of the proteasome from the 19S base or the 20S core do not induce a similar monopolar phenotype upon depletion. These results strengthen the growing belief that the proteasome may function not only as a 26S entity, but also as alternative or partial complexes that may each have their own independent cellular roles [15–21, 32, 60].

Our results uncover a novel role for the proteasome in regulating mitotic bipolar spindle formation. Specifically, we find that decreases in the 26:20S proteasome ratio downregulate the levels of the kinesin-5 motor protein, KIF11. As a primary factor in bipolar spindle assembly, even small changes in KIF11 abundance can have drastic effects on cellular growth and viability. For example, cells are extremely sensitive to minute amounts of KIF11 chemical inhibition (Fig. 3B) and the KIF11 gene is haploinsufficient [61]. Conversely, upregulation of KIF11 is also detrimental, as it promotes the occurrence of lagging chromosomes, centrosome fragmentation, micronuclei formation, and overall chromosome instability [62]. Thus, maintaining appropriate levels of KIF11 is essential for the survival and proper division of a cell, and the direct control of proteasome component abundances by the cell could be an important regulatory mechanism for preventing the lack or excess of KIF11 and proper spindle formation even in normal physiological conditions.

The proteasome 19S particle has important functions in substrate unfolding, ubiquitin recognition, and deubiquitination [3, 6–8]. Our data also suggests an additional role for the 19S particle as a negative regulator of proteasome degradation activity by the 20S core. We demonstrate that, in the absence of the 19S lid, degradation by the 20S particle can have deleterious consequences for the cell. Thus, the 19S lid plays a role not only in mediating ubiquitin-dependent degradation by the 26S proteasome, but also in preventing ubiquitin-independent degradation of a subset of substrates by the 20S core.

Negative regulation by the 19S lid is also important in the context of cell cycle progression. The role of ubiquitin-mediated degradation in regulation of the cell cycle is well-studied [9–11], but the effects of ubiquitin-independent degradation on mitotic progression have not been previously tested. The knockout of 19S components allowed us to analyze the cellular phenotypes associated with ubiquitin-independent degradation in mitotic cells. We reveal the adverse effects of ubiquitin-independent degradation on cell cycle progression and identify KIF11 as a distinct protein substrate of the 20S core. These results add to the current understanding of proteasomal function in cell cycle regulation and show that smooth progression through the cell cycle relies both on ubiquitin-dependent degradation from the 26S complex and on alternative modes of proteasome function.

In unperturbed cells, only half of the proteasome complexes are found in lidded forms, with the other half existing as uncapped 20S complexes [63, 64]. Despite this, the occurrence of monopolar spindles in a normal cell population is rare. How normal cells prevent the ectopic degradation of KIF11 by free 20S cores is still an open question and hints towards either an alternative negative regulator of 20S activity in normal cells or the existence of a regulatory component whose association upon loss of the 19S lid could control proteasomal substrate recognition. On the other hand, spindle formation is very sensitive to KIF11 levels [61, 62], thus it is possible that small changes in KIF11 levels induced by increased amounts of free 20S in PSMD-depleted cells could result in large differences in the percentage of properly formed spindles.

Finally, we found that PSMD-depleted cells arrest strongly in mitosis. These results suggest that PSMD subunits may be fruitful targets for cancer therapeutics, as cancerous cells are highly proliferative and would be susceptible to selective death through a mitotic arrest mechanism. Indeed, certain cancer cell types are reliant on high 26:20S ratios for their continued proliferation and high PSMD1 levels are correlated to poor survival prognosis in patients with oropharyngeal squamous cell carcinoma [65, 66]. Thus, the development of therapeutic strategies that either deplete 19S lid subunits or dissociate proteasome complexes to decrease the 26:20S ratio could lead to selective arrest and killing of highly proliferative cancer cells and improve patient outcomes. In addition, multiple myeloma cells are highly sensitive to proteasome inhibition. However, multiple myeloma patients eventually develop resistance to treatment with 20S proteasome inhibitors, such as bortezomib and carfilzomib [3, 67]. Recent work found that resistance in these cancerous cells is accompanied by a downregulation of 19S proteasome components and that the genetic knockdown of 19S subunits rendered multiple myeloma cells resistant to carfilzomib [67]. Here, we observed that the depletion of 19S proteasome lid subunits sensitized cells to chemical inhibition of KIF11 by *S*-Trityl-L-cysteine (STLC) treatment. Our results suggest that, in the face of multiple myeloma cell resistance to 20S inhibitors, a combination treatment of bortezomib/carfilzomib with STLC could selectively kill resistant cancerous cells through a monopolarity-induced mitotic arrest mechanism.

## Acknowledgments

We thank Gunter Sissoko for feedback on the manuscript, and members of the Cheeseman lab for feedback throughout the process. We would also like to thank members of the Whitehead Flow Cytometry and Proteomics Core Facilities. This work was supported by grants from the NIH/NIGMS (R35GM126930) and Chan-Zuckerberg Initiative (grant: Cell Biology @Scale) to IMC.

## Methods

### Cell culture and reagents

HeLa, A549, and 3T3 cell lines were cultured in Dulbecco’s modified Eagle medium (DMEM) supplemented with 10% fetal bovine serum (FBS), 100 U/ml penicillin and streptomycin, and 2mM L-glutamine at 37°C with 5% CO_2_. Doxycycline-inducible cell lines were cultured in medium containing tetracycline-free FBS. For CRISPR-Cas9 inducible knockout cell lines, *sp*Cas9 expression was induced with 1 µg/ml doxycycline hyclate (Sigma-Aldrich). Doxycycline media was renewed every 24 hours. Other drugs used on human cells were cycloheximide (50 µg/ml; Sigma-Aldrich), MG-132 (concentrations denoted in figures or figure legends; Enzo Life Sciences), taxol (1 µM; Invitrogen), thymidine (2 mM, Sigma-Aldrich), STLC (10 µM, unless otherwise noted in figure or figure legends; Sigma-Aldrich), and centrinone (200 nM, Oegema-Desai lab [69]). Transfections of Ub-R-GFP (Addgene Plasmid #11939) were done using Effectene Transfection Reagent (Qiagen).

### Cell Line Generation

Inducible CRISPR-Cas9 HeLa and H2B-mCherry HeLa cell lines were generated from previously described parental cell lines [33]. sgRNAs were cloned into the sgOpti plasmid (puro-resistant, Addgene plasmid #85681) and introduced into the parental cell line by lentiviral transduction, followed by selection with 0.5 µg/ml puromycin (Gibco). sgRNA guide sequences can be found in Supplementary Table 4.

GFP-KIF15 cell lines were generated by transfection using X-tremeGENE-9 (Roche) of GFP-KIF15 plasmid (gift from Patricia Wadsworth, [70]) followed by selection with 800 µg/ml Geneticin (G418 Sulfate, Thermo Fisher). Monoclonal cell lines were isolated by fluorescence-activated cell sorting and selected based on characteristic GFP localization by immunofluorescence.

GFP-KIF11 cell lines (including wild-type KIF11, KtoR KIF11, and KIF11 fragments) and the GFP control cell line were generated by lentiviral transduction. Lentivirus was made by transfection with Xtremegene-9 (Roche) of GFP or GFP-KIF11- containing lentivirus plasmid, VSV-G envelope plasmid, and psPAX2 (Addgene plasmid

#12260) packaging plasmid into HEK-293T cells. GFP-positive cells were then sorted by fluorescence-activated cell sorting.

### Western Blot

For western blot experiments, cells were harvested and flash frozen in liquid nitrogen to be stored at -80°C or immediately lysed on ice for 25 minutes in urea lysis buffer (50 mM Tris pH 7.5, 150 mM NaCl, 0.5% NP-40, 0.1% sodium dodecyl sulfate (SDS), 6.5 M Urea, 1X Complete EDTA-free protease inhibitor cocktail (Roche), 1 mM phenylmethylsulfonyl fluoride (PMSF)). Centrifugation was used to remove cellular debris. Protein concentrations were measured using Bradford reagent (Bio-Rad) to normalize loading. Lysates were mixed with Laemmli sample buffer and 2- mercaptoethanol and heated at 95°C for 5 minutes. Samples were loaded onto 8, 10, or 12% acrylamide gels for SDS-PAGE and transferred to nitrocellulose or Polyvinylidene Fluoride (PVDF) membrane (Cytiva). Membranes were blocked for 1 hour in blocking buffer (2.5% milk in Tris-Buffered Saline (TBS) + 0.1% Tween-20). Primary antibodies were diluted in 0.25% milk in TBS + 0.1% Tween-20 and added to the membrane for 2 hours at room temperature or overnight at 4°C. Membranes were washed with TBS + 0.1% Tween-20 and then incubated in either HRP-conjugated secondary antibodies (GE Healthcare; Digital) or in IRDye secondary antibodies (LI-COR) for one hour at room temperature. After washing in TBS + 0.1% Tween-20, membranes were either incubated in clarity-enhanced chemiluminescence substrate (Bio-Rad) and imaged using the KwikQuant Imager (Kindle Biosciences) or imaged using the Odyssey CLx Imager (LI-COR). Antibodies are listed in Supplementary Table 5.

### Immunofluorescence

For immunofluorescence experiments, cells were plated on poly-L-lysine (Sigma-Aldrich)-coated coverslips. Following fixation with either Formaldehyde or Methanol, coverslips were washed with phosphate-buffered saline (PBS) + 0.1% TritonX-100 (PBS-Tx) and blocked in blocking buffer (20 mM Tris-HCl, 150 mM NaCl, 0.1% Triton X- 100, 3% bovine serum albumin, 0.1% NaN_3_, pH 7.5) for 30 minutes or overnight. Primary antibodies were diluted in blocking buffer and added to coverslips for 1 hour or overnight. Cells were washed with 0.1% PBS-Tx. Cy2-, Cy3-, or Cy5-conjugated secondary antibodies (Jackson ImmunoResearch Laboratories) were diluted in blocking buffer and added to coverslips for 1 hour. Cells were washed with 0.1% PBS-Tx. 1 µg/ml Hoechst-33342 (Sigma-Aldrich) diluted in 0.1% PBS-Tx was added for 10 minutes to stain for DNA. Coverslips were mounted onto slides using ProLong^TM^ Gold antifade reagent (Invitrogen). Images were taken with a DeltaVision Ultra High-Resolution microscope and analyzed with Fiji (ImageJ, NIH). A list of antibodies and their dilutions can be found in Supplementary Table 5.

### Live-cell Imaging

For live-cell imaging, H2B-mCherry tagged cells were seeded onto 12-well glass-bottom plates (Cellvis, P12-1.5P). 1 hour prior to imaging, the media was changed to CO_2_- independent media (Gibco) supplemented with 10% FBS, 100 U/ml penicillin and streptomycin, and 2 mM L-glutamine, with or without 1 µM SiR-tubulin for visualizing microtubules. Imaging was conducted at 37°C on a DeltaVision Ultra High-Resolution microscope (40X) or with a Nikon eclipse microscope (20X). Images were analyzed with Fiji (ImageJ, NIH).

### Analysis of CRISPR-Cas9 cutting efficiency by TIDE

To analyze Cas9-based genome cutting efficiency, cell pellets from control and edited cells were resuspend in genomic DNA lysis buffer (100 mM Tris pH 8.0, 5 mM EDTA, 200 mM NaCl, 0.2% SDS, proteinase K to a final concentration of 0.2 mg/ml) and incubated overnight at 55**°**C. DNA was then precipitated the following day by addition of equal volume of isopropanol, then washed with 75% ethanol and resuspended in buffer (10 mM Tris-HCl pH 8.0, 0.1mM EDTA) overnight at 55°C in a shaking incubator. PCR was run using the isolated genomic DNA and the corresponding primers for each gene (Supplementary Table 4). PCR products were run on 1-2% agarose gels, gel extracted, and tested by Sanger sequencing (Azenta). Trace files were used for quantitative assessment of genome editing by TIDE (Tracking of Indels by Decomposition, https://tide.nki.nl/).

### siRNA treatment

Custom siRNAs against PSMD1 (pool: 5’-CAAAGGAUGCAGUACGGAA, 5’- AGACCAUACUGGAGUCGAA, 5’-GAUUGGAAGGCAUCGUAAA, 5’- CAUGGGAACUGCACGUCAA, or individual: 5’-AGACCAUACUGGAGUCGAA), PSMB7 (pool: 5’-CCGCAGGAAUGCCGUCUUG, 5’-GUAUCAAGGUUACAUUGGU, 5’- GGUCUAUAAGGAUGGCAUA, 5’- GAAGAUAAGUUUAGGCCAG), DYNC1H1 (pool: 5’-GAUCAAACAUGACGGAAUU, 5’-CAGAACAUCUCACCGGAUA, 5’- GAAAUCAACUUGCCAGAUA, 5’- GCAAGAAUGUCGCUAAAUU), and LIS1 (pool: 5’- CAAUUAAGGUGUGGGAUUA, 5’-UGAACUAAAUCGAGCUAUA, 5’- GGAGUGCCGUUGAUUGUGU, 5’- UGACAAGACCCUACGCGUA), and a non-targeting control pool (D-001206-13) were obtained from Dharmacon. siRNAs were used at a final concentration of 50 nM, unless otherwise stated in the text or figure legend. siRNAs and Lipofectamine RNAiMAX (Invitrogen) were mixed at equal volume in Opti-MEM Reduced Serum Medium (ThermoFisher), vortexed, and allowed to incubate for 20 minutes before adding to cells. For double siRNA experiments, final concentration of both siRNAs was 50 nM for both with double the volume of Lipofectamine RNAiMAX. Media was changed between 8-24 hours later back to full-serum media.

### RNA isolation, Reverse Transcription, and qPCR

To isolate RNA, 400 µl of TRIzol RNA isolation reagent (ThermoFisher) was added to cells in a 6-well plate. Lysed cells were then transferred to a 1.5 ml centrifuge tube, vortexed, and frozen at -80°C. After thawing samples, 120 µl of chloroform was added and tubes were vortexed vigorously for at least 30 seconds before centrifuging at 4°C. The aqueous phase was then transferred to a new tube with equal volume of chloroform, vortexed, and spun again at 4°C. Aqueous phase was transferred to a new tube with GlycoBlue Coprecipitant (Invitrogen), 5 M NaCl, and equal volume of isopropanol, and incubated on dry ice then centrifuged at 4°C. Pellets were washed with 75% Ethanol and resuspended in water. Reverse transcription was performed using Maxima First Strand cDNA Synthesis Kit for RT-qPCR (Thermo Scientific). For qPCR, cDNAs were diluted 1:20. 1 µM primers and 2X SYBR Green PCR Master Mix (Thermo Fischer

Scientific) were mixed with cDNAs in 384-well plates with three technical replicates per cDNA sample and primer pair. Primer sequences can be found in Supplementary Table 4.

### Immunoprecipitation

Beads were prepared prior to immunoprecipitation. NHS Mag Sepharose magnetic beads (Cytiva) were washed once with 500 µl of ice-cold 1 mM HCl, then mixed with 50 µg of GFP nanobody and Coupling Buffer B (0.2 M NaHCO_3_, 0.5 M NaCl, pH 8.3) to a total volume of 300 µl and incubated at room temperature for 1 hour. Beads were then washed with Blocking Buffer A (0.5 M ethanolamine, 0.5 M NaCl, pH 8.3) and Blocking Buffer B (0.1 M Sodium Acetate, 0.5 M NaCl, pH 4) and incubated in Blocking Buffer A for 15 minutes. After incubation, beads were washed again with Blocking Buffers A and B, then resuspended in 0.2 M ethanolamine, 0.2 M NaCl at 4°C.

Cells were harvested and resuspended in equal volume of 1X Lysis Buffer (50 mM HEPES, 1 mM EGTA, 1 mM MgCl_2_, 100 mM KCl, 10% glycerol) with 2% Triton X- 100, Complete EDTA-free protease inhibitor cocktail (Roche), 1 mM phenylmehtylsulfonyl fluoride (PMSF), 0.4 mM Sodium Orthovanadate, 5 mM Sodium Fluoride, 20 mM Beta-glycerophosphate, and ATP regeneration system (50 µg/ml creatine kinase, 35 mM creatine phosphate, 5 mM ATP). Cells were lysed on ice for 30 minutes prior to centrifuging at 15,000 rpm for 30 minutes. After centrifuging, ATP regeneration system was added to the supernatant before mixing with GFP-nanobody-conjugated beads for 2 hours at 4°C. After incubation, beads were washed with wash buffer (50 mM HEPES, 1 mM EGTA, 1 mM MgCl_2_, 100 mM KCl, 10% glycerol, 2% Triton X-100, ATP regeneration system, 1 mM DTT, 10 μg/mL leupeptin (Millipore), 10

μg/mL pepsin (Thermo Fisher Scientific), 10 μg/mL chymostatin (Millipore)), before elution with 0.1M Glycine pH 2.6. Eluted samples were mixed with 200 mM Tris pH 8.5 and 20% Trichloroacetic Acid (Fisher Bioreagents) overnight.

### Quantitative Mass Spectrometry

For whole-cell quantitative proteomics, cells were treated with either 50 nM PSMD1 siRNAs or 50 nM control siRNAs for 48 hours. For interphase cells, 2 mM Thymidine (Sigma-Aldrich) was added 24 hours prior to harvesting to arrest cells in S phase. For mitotic cells, 1 µM taxol (Invitrogen) was added 24 hours prior to harvesting to arrest cells in mitosis. Mitotic cells were isolated by mitotic shake off. Three biological replicates were collected for each condition. Cells were flash frozen in liquid nitrogen and stored at -80°C. To prepare protein extracts, cells were resuspended in RIPA buffer (225 mM NaCl, 1.5% Nonidet P-40 substitute, 0.75% Na Deoxycholate, 0.15% SDS, 75 mM Tris pH 8) with Complete EDTA-free protease inhibitor cocktail (Roche), 1 mM phenylmehtylsulfonyl fluoride (PMSF), 0.4 mM Sodium Orthovanadate, 5 mM Sodium Fluoride, and 20 mM Beta-glycerophosphate. Cells were lysed with Branson Digital Sonifier tip sonicator at 30% amplitude for 10 seconds once. Lysates were centrifuged at 20,000g for 30 minutes at 4°C. Supernatant was then mixed with 20% Trichloroacetic Acid (Fisher Bioreagents) overnight on ice to precipitate proteins. The next day, samples were centrifuged at 20,000g at 4°C. Pellets were washed two times with cold acetone and dried in an Eppendorf Vacufuge.

To prepare samples for mass spectrometry, pellets were first resuspended in SDS lysis buffer (5%, 50 mM TEAB pH 8.5) with 20 mM DTT and incubated at 95°C for 10 minutes. Samples were then incubated with 40 mM iodoacetamide (Sigma) for 30 minutes in the dark. 1.2% phosphoric acid was added to the sample, which was then run over S-Trap microcolumns (ProtiFi) and subjected to tryptic digest on the columns overnight (Promega). Digested peptides were then eluted, quantified with Quantitative Fluorometric Peptide Assay Kit (Pierce), and lyophilized. A more detailed protocol for S-Trap digestion and elution can be found in the ProtiFi S-trap micro kit protocol.

For TMT-based quantitative mass spectrometry, peptides from each sample were mixed with different TMT10plex labels at a 1:10 ratio by mass. Labeling occurred in 30% acetonitrile and 24.5 mM TEAB pH8.5 for 1 hour at room temperature. After 1 hour, the reaction was quenched with hydroxylamine for 15 minutes at room temperature. TMT-labeled samples were then pooled, lyophilized, and stored at -80°C. Next, samples were fractionated using Pierce High pH Reversed-Phase Peptide Fractionation kit (Pierce). Fractions were lyophilized and resuspended in 0.1% formic acid and analyzed by mass spectrometry with an Orbitrap Exploris 480. Protein identification and Tandem Mass Tag quantification was done in Proteome Discoverer 2.4 (Thermo Fisher Scientific).

### Native Gel and Suc-LLVY-AMC assay

Cells were harvested and resuspended in TSDG buffer (10 mM Tris HCl pH 8, 1.1 mM MgCl_2_, 10 mM NaCl, 0.1 mM EDTA, 1 mM NaN_3_, 1 mM DTT, 2 mM ATP, 10% glycerol). Lysis was carried out through 7 sequential freeze-thaws in liquid nitrogen. Lysates were centrifuged for 30 minutes at 4°C and supernatant was collected. Samples were mixed with glycerol and run on a 3.5% acrylamide gel containing 0.5mM ATP in Native Running Buffer (90mM Tris Borate, 5mM MgCl_2_, 0.5mM EDTA, 0.5mM ATP) for 3 hours at 100V.

For Suc-LLVY-AMC assay: Following running, native gel was incubated in Suc-LLVY-AMC reaction buffer (50mM Tris pH 7.5, 0.5 mM ATP, 0.5 mM DTT, 0.04 mM Suc-LLVY-AMC substrate (Fischer Scientific)) for 30 minutes at 37°C. For activation of 20S particles, gel was imaged in reaction buffer + 0.02% SDS for 30 minutes at 37°C. Gels were then imaged with ChemiDoc Imaging System (Bio-Rad).

---

**Extended Data Figure 1:**
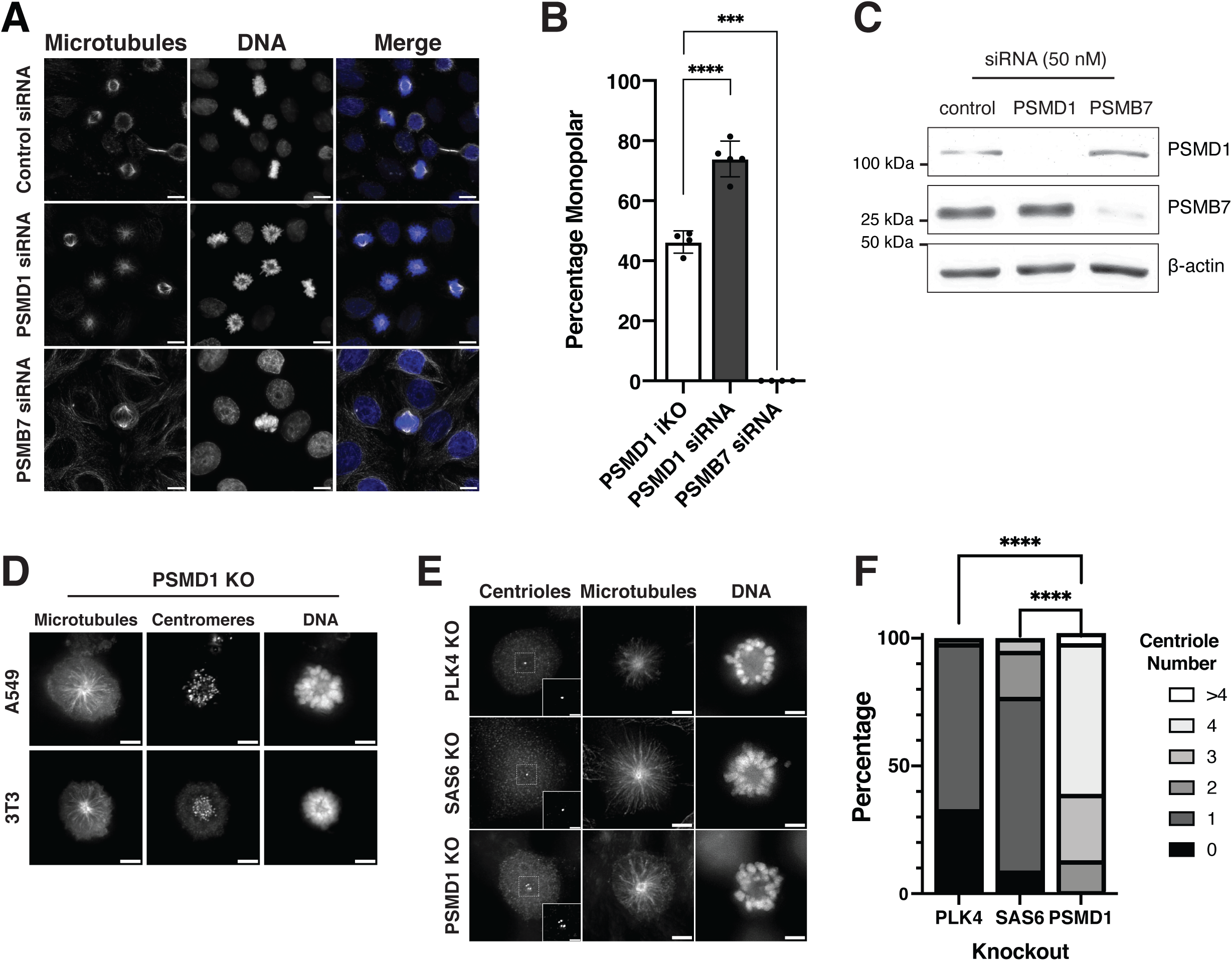
Alternative strategies for inducing proteasome subunit depletion and counting centriole numbers A. Representative immunofluorescence images of asynchronous HeLa cells treated with 50nM control siRNAs, HeLa cells treated with 50 nM PSMD1 siRNAs, and HeLa cells treated with 50 nM PSMB7 siRNAs. Scalebars = 10 µm B. Percentage of mitotic cells with monopolar spindles observed in inducible CRISPR-Cas9 PSMD1 knockout HeLa cells, HeLa cells treated with 50 nM PSMD1 siRNAs, and HeLa cells treated with 50 nM PSMB7 siRNAs. Bars represent mean ± standard deviation. P-values were calculated with two-tailed Welch’s t-tests: **** represents p< 0.0001, *** represents p = 0.0001. C. Western blot of cells treated with control, PSMD1, or PSMB7 siRNAs. Blot was incubated with PSMD1 or PSMB7 antibodies. β-actin is used as a loading control. D. Representative immunofluorescence images of monopolar mitotic A549 and 3T3 cells resulting from knockout of PSMD1. Cells were transduced with lentivirus containing mCherry-expressing gene knockout plasmids for PSMD1 and sorted for mCherry before seeding on coverslips. Scalebars = 5 µm. E. Representative immunofluorescence images of monopolar mitotic cells from inducible CRISPR-Cas9 knockouts of PSMD1, PLK4, and SAS6. Cells were stained for centrioles with Centrin2 antibody. Regions enlarged in insets are outlined with dashed squares. Image brightness not identical. Scalebars for full-sized images = 5 µm. Scalebars for insets = 2 µm. F. Quantification of centriole number in inducible CRISPR-Cas9 knockouts of PSMD1, PLK4, and SAS6. Bars show the percentage of cells in each centriole number category. Experiment was replicated 3 times, 31-60 cells were quantified for each condition for each replicate. P-values were calculated with chi-square tests. **** represents p< 0.0001.

**Extended Data Figure 2.**
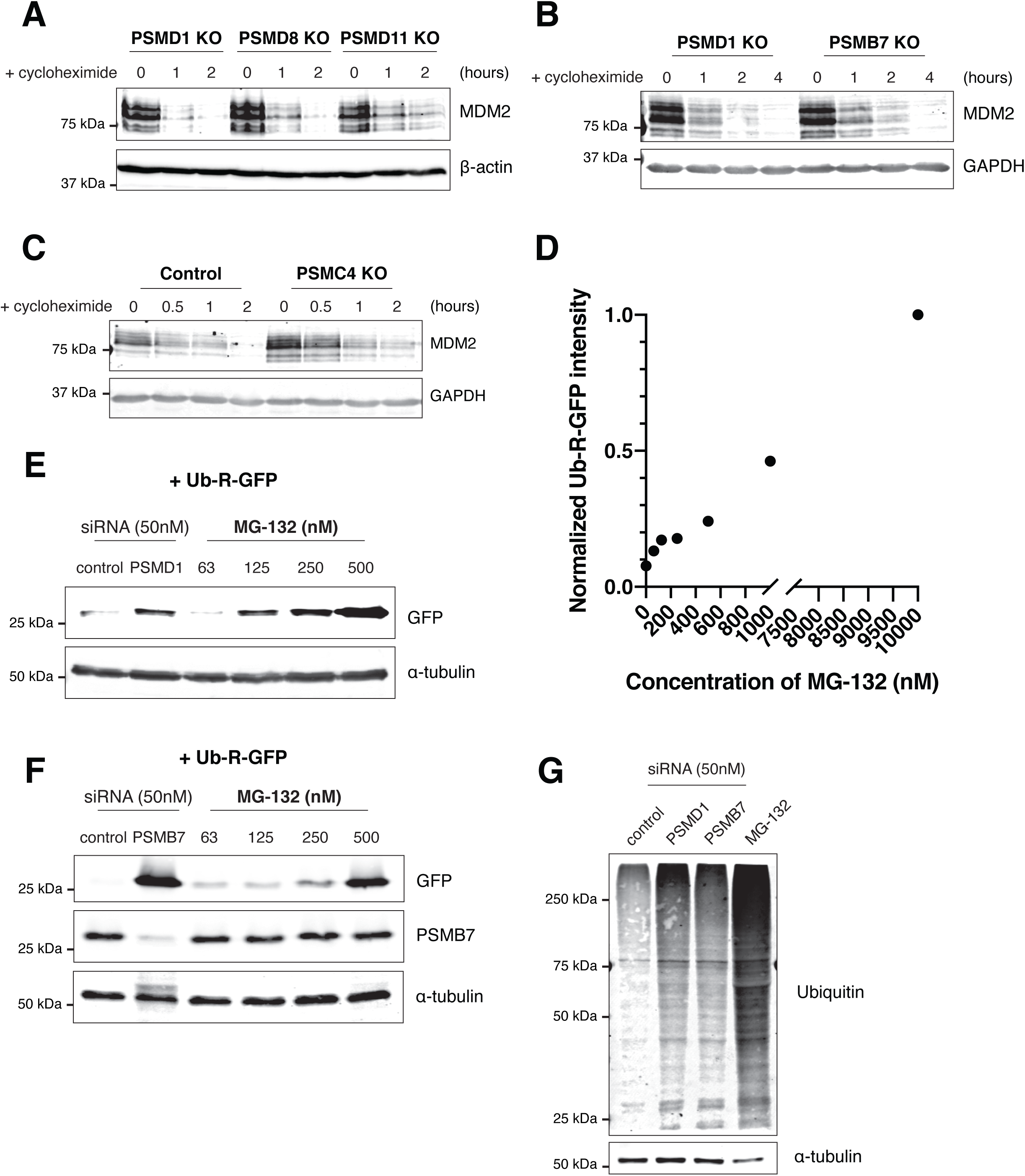
: Assays for ubiquitin-mediated proteasome degradation activity A. Western blot of inducible CRISPR-Cas9 PSMD1, PSMD8, and PSMD11 knockout cells treated with 50 µg/ml of cycloheximide for the indicated amount of time. Blot was incubated with MDM2 antibody. β-actin is used as a loading control. B. Western blot of inducible CRISPR-Cas9 PSMD1 and PSMB7 knockout cells treated with 50 µg/ml of cycloheximide for the indicated amount of time. Blot was incubated with MDM2 antibody. GAPDH is used as a loading control. C. Western blot of control and inducible CRISPR-Cas9 PSMC4 knockout cells treated with 50 µg/ml of cycloheximide for the indicated amount of time. Blot was incubated with MDM2 antibody. GAPDH is used as a loading control. D. Graph quantifying (Figure 2C). GFP bands were quantified and divided by the corresponding intensity of β-actin bands. E. Western blot of Ub-R-GFP-transfected cells treated with different conditions: 50 nM control siRNAs, 50 nM PSMD1 siRNAs, and the indicated concentrations of MG-132. Cells were transfected with Ub-R-GFP plasmid 60 hours prior to collection. siRNAs were added 48 hours prior to collection, and MG-132 was added 24 hours prior to collection. Blot was incubated with GFP antibody. β- actin is used as a loading control. F. Western blot of Ub-R-GFP-transfected cells treated with different conditions: 50 nM control siRNAs, 50 nM PSMB7 siRNAs, and the indicated concentrations of MG-132. siRNAs were added 72 hours prior to collection. Cells were transfected with Ub-R-GFP plasmid 48 hours prior to collection, and MG-132 was added 24 hours prior to collection. Blot was incubated with GFP antibody and PSMB7 antibody. α-tubulin is used as a loading control. G. Western blot of cells treated with 50 nM control siRNAs, 50 nM PSMD1 siRNAs, 50 nM PSMB7 siRNAs, or 10 µM MG-132. Control and PSMB7 siRNAs were added 66 hours prior to collection, PSMD1 siRNAs were added 48 hours prior to collection, and MG-132 was added 24 hours prior to collection. Blot was incubated with Ubiquitin antibody. α-tubulin is used as a loading control.

**Extended Data Figure 3:**
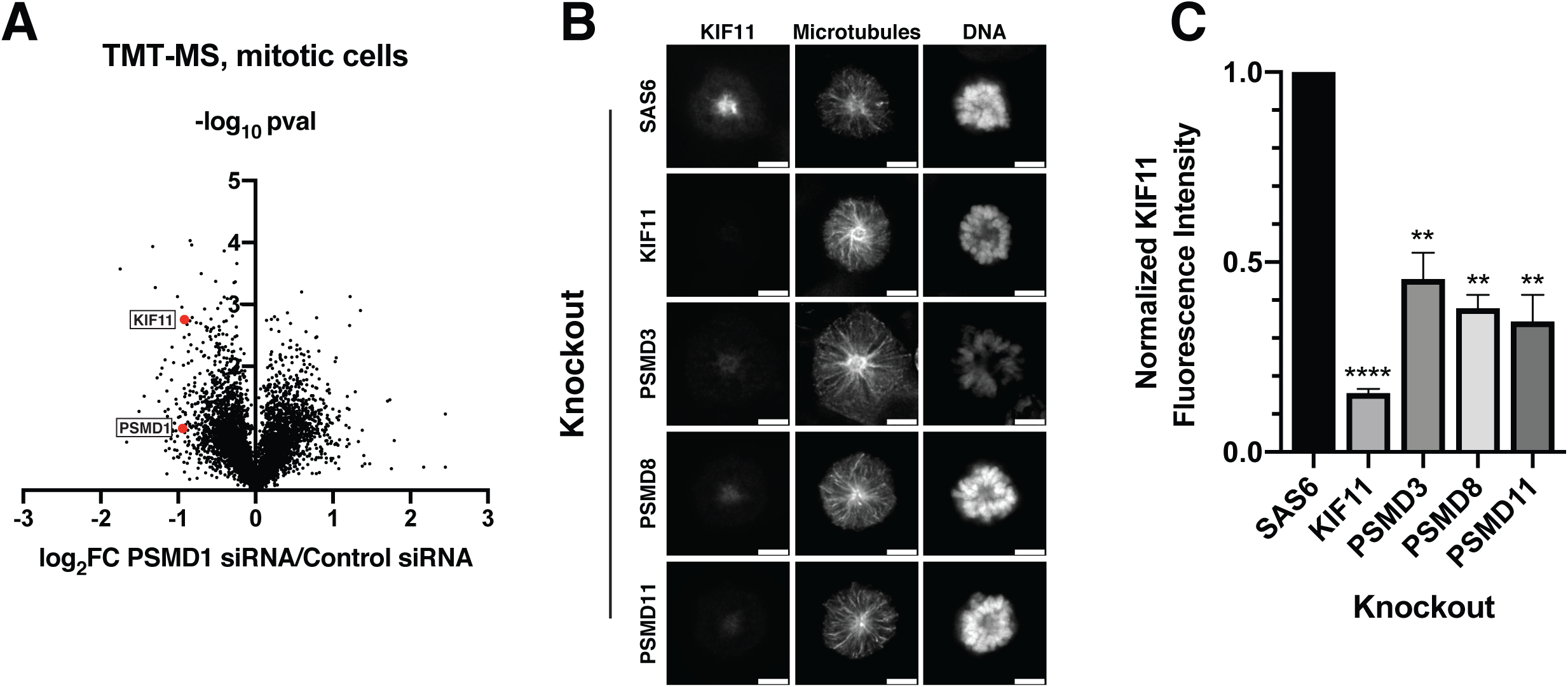
KIF11 is lost from mitotic cells upon PSMD1 depletion A. Comparison of protein abundances in cells treated with 50 nM PSMD1 siRNAs and cells treated with 50 nM control siRNAs. Cells were treated with siRNAs for 48 hours and arrested in mitosis with 1 µM taxol prior to harvesting. Mitotic cells were specifically isolated from interphase cells using mitotic shake off. Protein abundances were obtained using TMT-based (Tandem Mass Tag) quantitative mass spectrometry of three replicates for PSMD1 RNAi condition and two replicates for control RNAi condition. Volcano plot shows log_2_ of protein abundances in PSMD1 knockdown cells divided by protein abundances in control knockdown cells vs. -log_10_ of p-values. Values on the right of the plot represent proteins with increased abundance in PSMD1 knockdown cells and values on the left represent proteins with decreased abundance in PSMD1 knockdown cells. PSMD1 and KIF11 are highlighted in red. B. Representative immunofluorescent images of monopolar mitotic cells showing KIF11 levels in inducible CRISPR-Cas9 knockouts of SAS6, KIF11, PSMD3, PSMD8, and PSMD11. Scalebar = 5 µm. C. Quantification of KIF11 fluorescence intensity in monopolar mitotic cells in different conditions. Fluorescence intensity values were normalized to levels in control SAS6 knockout cells. Bars represent mean ± standard deviation. P-values were calculated with two-tailed Welch’s t-tests comparing each condition to SAS6 knockout: p< 0.0001 for KIF11, p = 0.0053 for PSMD3, p = 0.0011 for PSMD8, and p = 0.0037 for PSMD11. **** represents p < 0.0001, ** represents p < 0.01. Experiment was replicated 3 times, 38-76 cells were quantified for each condition for each replicate.

**Extended Data Figure 4:**
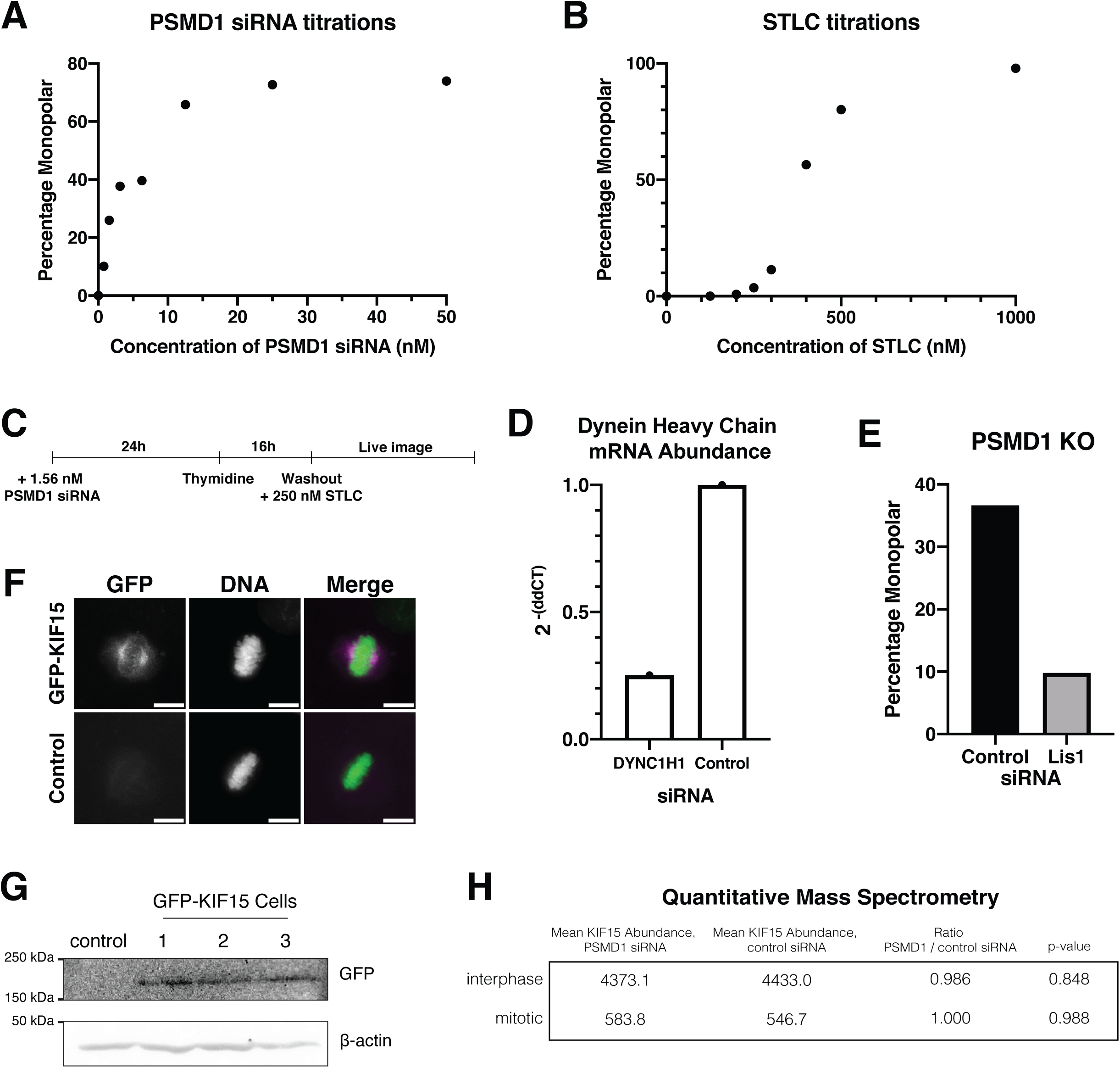
Forces acting on the mitotic spindle can be experimentally manipulated A. Titration curve for PSMD1 siRNA titrations. PSMD1 siRNA concentrations were between 0 and 50 nM. The percentage of mitotic cells with monopolar spindles was quantified by immunofluorescence. B. Titration curve for STLC titrations. STLC concentrations ranged from 0 to 1000 nM. The percentage of mitotic cells with monopolar spindles was quantified by immunofluorescence. C. Diagram showing the experimental timeline for the STLC/PSMD1 siRNA synergy experiment shown in Fig. 4A. D. Graph showing the abundance of dynein heavy chain (DYNC1H1) mRNA as quantified by qPCR following treatment of cells with 50 nM of DYNC1H1 or control siRNAs. 3 technical replicates were conducted per condition for a single biological replicate. CT values for DYNC1H1 mRNA were normalized to those of GAPDH before normalization to control siRNA condition. E. Quantification of live imaging experiment. Bar graph showing the percentage of cells entering mitosis with monopolar spindles in PSMD1 knockout cells treated with either 50 nM of control or Lis1 siRNA. Experiment was conducted 1 time and between 1300-1406 mitotic cells were counted for each condition. F. Representative immunofluorescence images of GFP-KIF15 and control cells. Cells were stained with GFP booster. GFP-KIF15 localizes to the spindles. Scalebar = 10 µm G. Western blot of GFP-KIF15 monoclonal cells lines and control parental cells blotted with GFP antibody. Size of KIF15 is 160 kDa. β-actin is used as a loading control. H. Table showing abundance of KIF15 in PSMD1 siRNA- vs. control siRNA-treated cells. Protein abundances were obtained using TMT-based (Tandem Mass Tag) quantitative mass spectrometry.

**Extended Data Figure 5:**
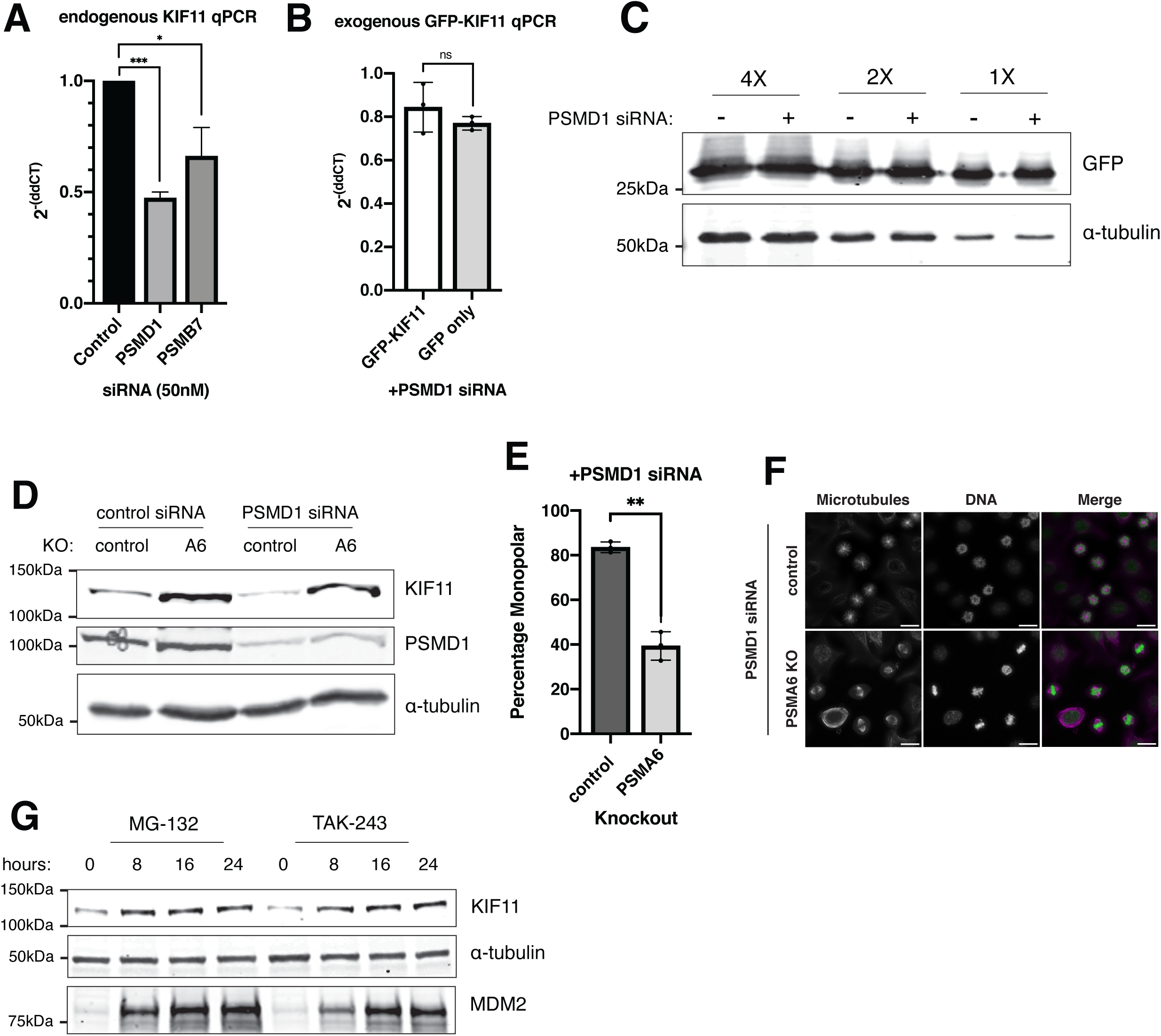
KIF11 is degraded by the 20S proteasome in PSMD1- depleted cells A. Graph showing the abundance of endogenous KIF11 mRNA in different conditions as quantified by qPCR following treatment of cells with 50 nM control, PSMD1, or PSMB7 siRNAs. 3 technical replicates are shown per condition for a single biological replicate. Experiment was replicated two times with similar results. CT values for KIF11 mRNA were normalized to those of GAPDH before normalization to control siRNA condition. B. Graph showing the relative decrease in the abundances for the indicated mRNAs following treatment of cell lines with PSMD1 siRNAs. GFP-KIF11 or GFP only cell lines were treated with 50 nM PSMD1 siRNAs for 48 hours prior to harvesting. 3 biological replicates with 3 averaged technical replicates each are shown per condition. CT values for KIF11 mRNA or GFP mRNA were normalized to those of GAPDH before normalization to respective control siRNA condition. C. Western blot of control GFP-expressing cells treated with 50 nM control (-) or PSMD1 (+) siRNAs. Blot was incubated in GFP antibody. The same samples were loaded at different concentrations (4X, 2X, or 1X). α-tubulin is used as a loading control. D. Western blot of control cells or PSMA6 knockout cells treated with either 50 nM control or PSMD1 siRNAs. Knockouts were generated by transducing HeLa cells with lentivirus containing the mCherry-expressing gene knockout plasmids for PSMA6. Cells were then sorted for mCherry positive cells by FACS. Blot was incubated with KIF11 and PSMD1 antibodies. α-tubulin is used as a loading control. E. Quantification of immunofluorescence experiment from PSMA6 knockout cells (Extended Data Figure 5D). Bar graph shows the percentage of mitotic cells with monopolar spindles for control or PSMA6 knockout cells treated with 12.5 nM PSMD1. P-value was calculated with two-tailed Welch’s t-tests, p = 0.0031. ** represents p < 0.01. Experiment was conducted 3 times. F. Representative immunofluorescence images from Extended Data Fig. 5E. Control (top) or PSMA6 knockout cells (bottom) were treated with 12.5 nM PSMD1 siRNAs. Scalebar = 20 µm. G. Western blot of control cells treated with MG-132 or TAK-243 for the indicated amount of time. Blots were incubated with KIF11 or MDM2 antibodies. α-tubulin is used as a loading control. KIF11 Separate blots were used with the same samples and same amounts loaded. One blot was incubated with KIF11 and α- tubulin antibodies and the other with MDM2 antibody.

**Extended Data Figure 6:**
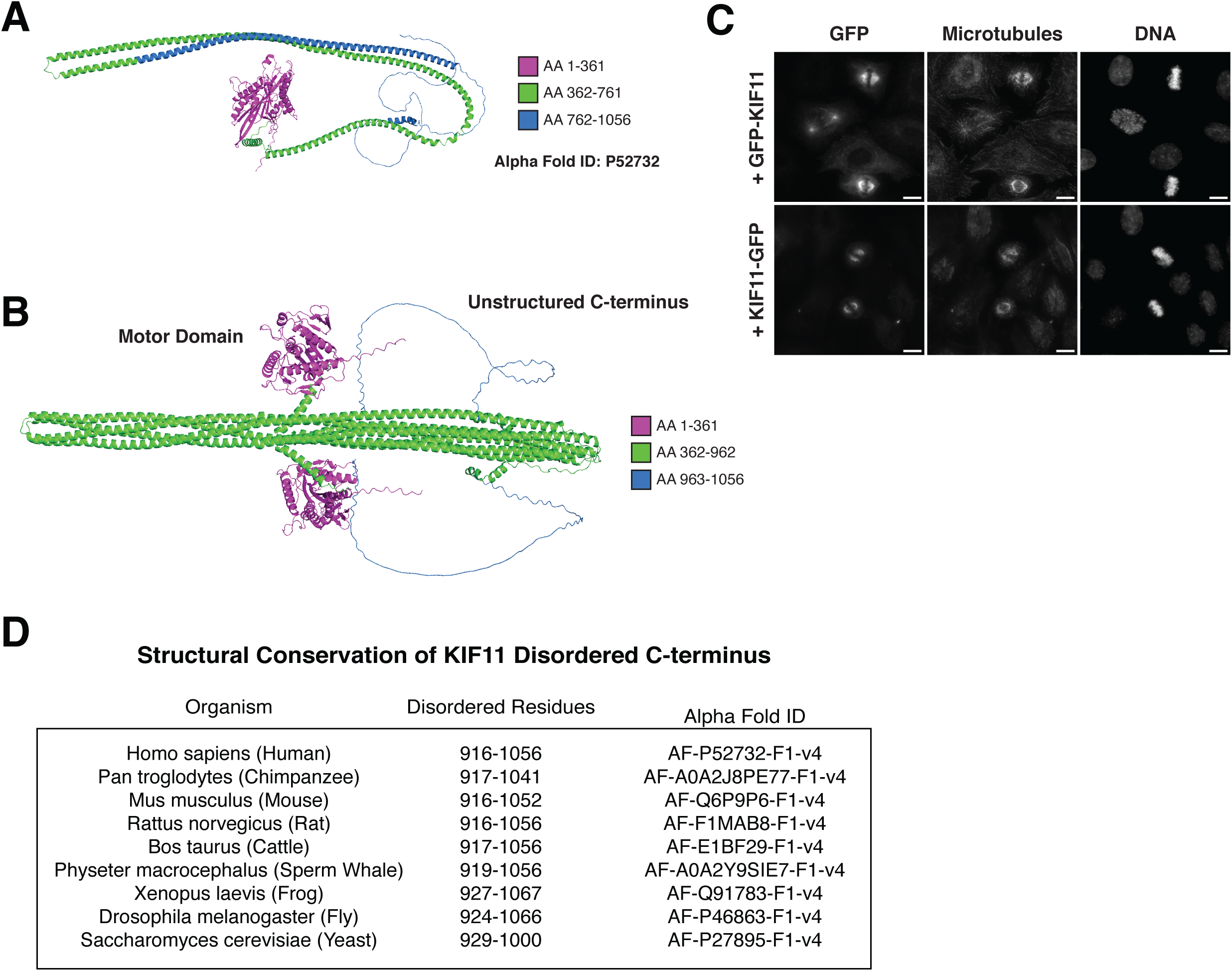
KIF11 has a conserved unstructured C-terminus A. Alpha-fold prediction of full-length Homo sapiens KIF11 structure. AF-P52732- F1-v4. B. Alpha-fold prediction of full-length dimer of Homo sapiens KIF11 structure. C. Representative images of GFP-KIF11-expressing cell lines. Cell lines shown characteristic KIF11 localization at the spindles. Scale bars = 10 µm. D. Table showing the conservation of KIF11 unstructured C-terminus across different species.

**Supplementary Table 1:** TMT-based quantitative mass spectrometry data from interphase PSMD1 RNAi and control RNAi cells. Cells were treated with either 50 nM PSMD1 siRNAs or 50 nM control siRNAs for 48 hours and then arrested in S phase with 2 mM thymidine prior to harvesting. Three replicates for each condition were conducted. Protein abundances for each replicate are shown, as well as the mean of triplicate abundances for each condition, the abundance ratio of PSMD1 RNAi/control RNAi, and associated p-values.

**Supplementary Table 2:** TMT-based quantitative mass spectrometry data from mitotic PSMD1 RNAi and control RNAi cells. Cells were treated with either 50 nM PSMD1 siRNAs or 50 nM control siRNAs for 48 hours and then arrested in mitosis with 1 µM taxol prior to harvesting. Mitotic cells were specifically isolated from interphase cells using mitotic shake off. Three replicates for PSMD1 RNAi condition and two replicates for control RNAi condition were conducted. Protein abundances for each replicate are shown, as well as the mean of replicate abundances for each condition, the abundance ratio of PSMD1 RNAi/control RNAi, and associated p-values.

**Supplementary Table 3:** TMT-based quantitative mass spectrometry data from GFP immunoprecipitations of GFP-PSMB4 cells treated with PSMD1 RNAi or control RNAi cells. GFP-PSMB4 cells were treated with either 50 nM PSMD1 siRNAs or 50 nM control siRNAs for 48 hours and with 2 mM thymidine prior to harvesting. Three replicates for PSMD1 RNAi condition and two replicates for control RNAi condition were conducted. Protein abundances for each replicate are shown, as well as the mean of replicate abundances for each condition, the abundance ratio of PSMD1 RNAi/control RNAi, and associated p-values.

**Supplementary Table 4:** Oligonucleotide sequences. *Top*: CRISPR-Cas9 knockout guide sequences. *Middle*: Primer sequences used to amplify genomic DNA fragments by PCR from knockout cells for analysis of genome cutting efficiency by TIDE (Tracking of Indels by Decomposition). *Bottom*: Primer sequences used for qPCR.

**Supplementary Table 5:** List of antibodies used and their dilutions.

## Notes

### Competing Interest Statement

The authors have declared no competing interest.

