## Supplementary material for "19S proteasome loss causes monopolar spindles through ubiquitin-independent KIF11 degradation": Tables

**Table 1: Percentage editing of inducible CRISPR-Cas9 knockouts.** Genome cutting efficiency of proteasome subunit CRISPR-Cas9 inducible knockouts as measured by TIDE (Tracking of Indels by Decomposition).

| **Complex** | **Subunit** | **Percentage Editing** |
| --- | --- | --- |
| 19S Base | PSMD1 | 40.5 |
|  | PSMD2 | 56.3 |
|  | PSMC2 | 48.6 |
|  | PSMC4 | 49.1 |
|  | PSMC6 | 29.0 |
| 19S Lid | PSMD3 | 34.3 |
|  | PSMD6 | 20.3 |
|  | PSMD7 | 27.4 |
|  | PSMD8 | 49.0 |
|  | PSMD11 | 44.3 |
|  | PSMD12 | 40.8 |
|  | PSMD13 | 27.5 |
|  | PSMD14 | 49.2 |
|  | ADRM1 | 55.4 |
| 20S Core | PSMA1 | 46.3 |
|  | PSMA6 | 46.3 |
|  | PSMB1 | 39.6 |
|  | PSMB5 | 49.1 |
|  | PSMB7 | 40.7 |

**Table 2: Percentage editing of “one-shot” CRISPR-Cas9 knockouts.** Genome cutting efficiency of proteasome subunit CRISPR-Cas9 knockouts generated by lentiviral transduction of mCherry-containing gene knockout plasmids, as measured by TIDE (Tracking of Indels by Decomposition).

| **Subunit** | **Percentage Editing** |
| --- | --- |
| PSMD1 | 84.7 |
| PSMD8 | 78.6 |
| PSMD11 | 94.0 |
| PSMA6 | 65.6 |
| PSMC4 | 52.6 |
